# Phylogenomics of neglected flagellated protists supports a revised eukaryotic tree of life

**DOI:** 10.1101/2024.05.15.594285

**Authors:** Guifré Torruella, Luis Javier Galindo, David Moreira, Purificación López-García

## Abstract

Eukaryotes evolved from prokaryotic predecessors in the early Proterozoic^1,2^ and radiated from their already complex last common ancestor^3^, diversifying into several supergroups with unresolved deep evolutionary connections^4^. They evolved extremely diverse lifestyles, playing crucial roles in the carbon cycle^5,6^. Heterotrophic flagellates are arguably the most diverse eukaryotes^4,7-9^ and often occupy basal positions in phylogenetic trees. However, many of them remain undersampled^4,10^ and/or *incertae sedis*^4,11-18^. Progressive improvement of phylogenomic methods and a wider protist sampling have reshaped and consolidated major clades in the eukaryotic tree^13-19^. This is illustrated by the Opimoda^14^, one of the largest eukaryotic supergroups (Amoebozoa, Ancyromonadida, Apusomonadida, Breviatea, CRuMs, Malawimonadida, and Opisthokonta –including animals and fungi–)^4,14,19-22^. However, their deepest evolutionary relationships still remain uncertain. Here, we sequenced transcriptomes of poorly studied flagellates^23,24^ (fourteen apusomonads^25,26^, seven ancyromonads^27^ and one cultured Mediterranean strain of *Meteora sporadica*^17^) and conducted comprehensive phylogenomics analyses with an expanded taxon sampling of early-branching protists. Our findings support the monophyly of Opimoda, with CRuMs being sister to the Amorphea (amoebozoans, breviates, apusomonads, and opisthokonts), and ancyromonads and malawimonads forming a moderately supported clade. By mapping key complex phenotypic traits onto this phylogenetic framework, we infer an opimodan biflagellate ancestor with an excavate-like feeding groove, which ancyromonads subsequently lost. While breviates and apusomonads retained the ancestral biflagellate state, some early-diverging Amorphea lost one or both flagella, facilitating the evolution of amoeboid morphologies, novel feeding modes, and palintomic cell division resulting in multinucleated cells. These innovations likely facilitated the subsequent evolution of fungal and metazoan multicellularity.

## Results and Discussion

To contribute solving the deepest evolutionary relationships among eukaryotic taxa, we sought to enrich the available taxon sampling in early diverging lineages of flagellates, many of which are understudied grazers dwelling on benthos^6,13,15,18,25^. Although molecular environmental approaches reveal a wide diversity of such lineages^24^, the most complete genome/transcriptome data are generally retrieved from cultured protists^28^. Accordingly, we took advantage of our recent cultivation of several predatory flagellates sampled from marine and freshwater sediments, including several opisthokont-related^22^ apusomonads^26^, and *incertae sedis* ancyromonads^27^. We also studied a stably cultured Mediterranean strain of the long enigmatic *Meteora sporadica*^17^ (Figure S1), which showed some early affinity with the Diphoretickes^12,13^, more recently confirmed^18^. Diaphoretickes, together with the Opimoda, are considered one of the largest eukaryotic supergroups^19,29^.

### High-quality curated transcriptomes of deep-branching flagellates

We generated transcriptomic datasets for Apusomonadida^23^ and Ancyromonadida^11^ species by sequencing polyA-containing transcripts of recently cultured species spanning the diversity of these two clades^26,27^. We also sequenced the transcriptome of *Meteora sporadica*^17^ CRO19MET, isolated from the same Mediterranean area as the originally described type species^30^ (Table S1). Since these species were in co-culture with diverse bacteria and sometimes other protists, we established a pipeline (Figure S2) to eliminate contaminant sequences (see Methods; Tables S2-S4). After strict manual curation, we obtained highly complete transcriptomes (most more than 80% and up to ∼93% complete), as assessed by the presence of universal single-copy genes using BUSCO^31^ (Table S1). Since de novo transcriptomes often show artificially duplicated sequences (artefactual isoforms generated by assembly programs), we tested inferred oligopeptide redundancy by clustering sequences with CD-HIT at 90% identity. This procedure removed few such oligopeptide sequences for most species (6.5% on average), except for the highly duplicated *Mylnikovia oxoniensis* (∼42%), *Multimonas media* (20.5%), *Apusomonas australiensis* (15.5%) and *Cavaliersmithia chaoae* (9.6%). The removal of this redundancy did not affect the BUSCO completeness (Table S2).

### Phylogenomic analyses of an expanded dataset of heterotrophic flagellates

To infer the evolutionary relationships among major eukaryotic supergroups, we searched for a collection of commonly used conserved phylogenetic markers^13^ in our raw transcriptome data. Based on individual phylogenetic trees, we eliminated non-orthologous sequences (see Methods), which resulted in a curated dataset of 303 markers for 101 taxa, representing 97,171 amino acid positions, for downstream phylogenomic analyses. This marker set displayed few missing data across most of our species in comparison with previously sequenced apusomonad and ancyromonads transcriptomes (Table S5). We applied state-of-the-art complex mixture models of sequence evolution in both maximum likelihood (ML) and Bayesian inference (BI) methods to alleviate putative homoplasy and long-branch attraction artefacts. We reconstructed an ML tree using the LG+C60+G4 model with 1,000 ultrafast bootstraps (ufbs; Figure 1 and S3A), and used it as a guide tree for the posterior mean site frequencies (PMSF) approximation to calculate non-parametric bootstrap support (npbs; 100 replicates; Figure 1). We also reconstructed an ML tree using the eukaryote-linked model (ELM) of sequence evolution^32^ under ELM+C60+G4 (1,000 ufbs; Figure S3C) to guide a PSMF approximation and calculate npbs (100 replicates; Figure S3B). For BI analyses, we used two MCMC chains with the CAT-GTR (Figure S3D) and CAT-Poisson (Figure S3E) models of sequence evolution. BI and ML phylogenomic analyses yielded congruent tree topologies for major eukaryotic supergroups, although there were a few changes in the position of some branches (Figure 1 and Figure S3; see below).

**Figure 1.**
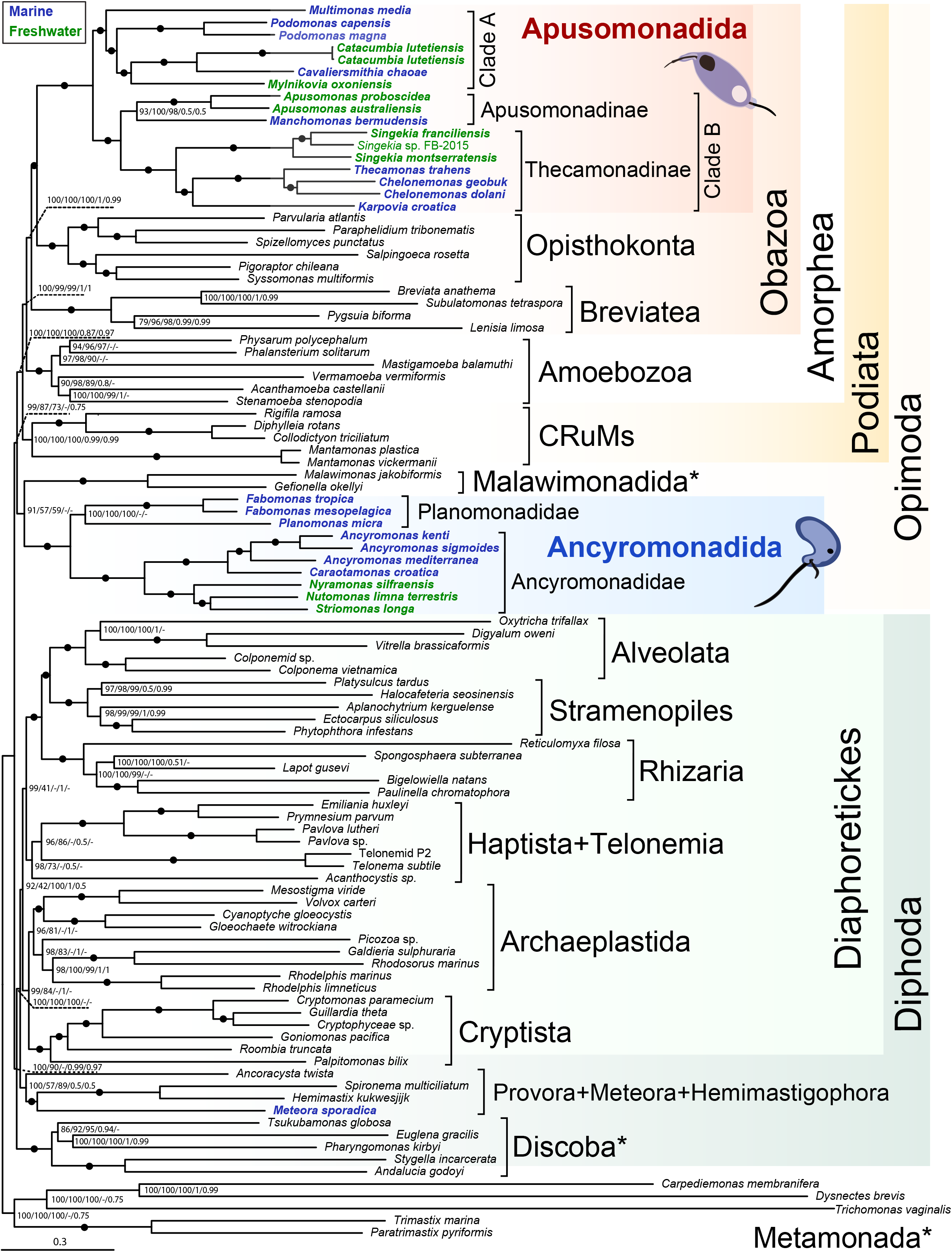
Phylogenomic tree of eukaryotes including an expanded diversity of Apusomonadida and Ancyromonadida. The tree was inferred using Maximum Likelihood from 97,171 amino acid positions and 101 taxa (model of sequence evolution: LG+C60+G4). The root of the tree has been arbitrarily placed between Metamonada and the rest of eukaryotes. Numbers at nodes represent non-parametric bootstrap (npbs) percentages under the PMSF approach (100 replicates); ultrafast bootstrap percentages of the ML LG+C60+G4 tree (1,000 replicates); npbs under the PMSF approach of the ML ELM+C60+G4 model tree (100 replicates); and Bayesian posterior probabilities under CAT-GTR and CAT-Poisson models, respectively. Black dots denote maximum support with all methods. The scale bar indicates the number of expected substitutions per unit branch length. The asterisks indicate taxa of typical excavates.

All our analyses retrieved the monophyly of Apusomonadida and Ancyromonadida with maximal support. The internal relationships within each of these groups were overall stable, except for minor uncertainties (Figure 1). The Apusomonadida comprised two major clades, A and B, grouping a mix of marine and freshwater species showing an overall internal topology congruent with 18S rRNA gene phylogenies^26^. Clade A grouped species with elongated *Amastigomonas*-type morphology and relative large cell size (∼8 µm; genera *Multimonas, Podomonas, Cavaliersmithia, Mylnikovia* and *Catacumbia*). Clade B comprised Thecamonadinae, grouping *Amastigomonas*-type phenotypes of smaller size (∼5 µm cell diameter; *Singekia, Karpovia, Chelonemonas* and *Thecamonas*)^26^, and Apusomonadinae, which grouped the genera *Apusomonas*, characterized by rounded cells, and *Manchomonas*, exhibiting elongated and larger cells (9.5 µm)^33^. The only internal uncertainty within the apusomonads concerned the support for the monophyly of *Manchomonas* and *Apusomonas*, very low in both BI trees (0.5 posterior probability –PP–; Figure 1 and S3D-E) in comparison with the high npbs and ufbs values in ML trees (93/98% LG+C60+G4/ELM+C60+G4 npbs; 100% LG+C60+G4 ufbs; Figures S3A-C). These differences in support may result from the *Manchomonas bermudensis* elevated missing data (90.49% gaps; only 62 markers retrieved from the available EST data –F. Lang, unpublished–; Table S5). Apusomonadida formed a fully supported clade with Opisthokonta, which was sister to the Breviatea (Obazoa)^14^. In turn, Obazoa were sister to Amoebozoa with full support (Amorphea) and Amorphea to CRuMs with very good to full support, forming the supergroup Podiata^34^ (Figure 1).

Similarly, we retrieved the monophyly of the two described ancyromonad families, Ancyromonadidae and Planomonadidae^35^. Their internal topology was congruent with that based on 18S rRNA genes^27^. Within Ancyromonadidae, we recovered a clade of marine (*Caraotamonas* and *Ancyromonas* species) and another of freshwater (*Striomonas, Nutomonas* and *Nyramonas*) representatives (Figure 1). However, we only retrieved full support for the *Planomonas* and *Fabomonas* (Planomonadidae) monophyly with ML; BI analyses placed *Fabomonas* as the earliest-branching ancyromonad (0.95 PP for CAT-GTR, and 0.99 PP for CAT-Poisson; Figure S3D-E). Interestingly, malawimonads^12^, traditionally classified within Excavata^36-38^, branched sister to the ancyromonads in our ML trees (Figure 1, Figure S3A-C), in line with some previous analyses^14,39^, although with relatively low support, especially for ELM analyses (59% npbs; 85% ufbs; Figure S3B-C). Ancyromonadida and Malawimonadida formed a monophyletic supergroup with Amorphea and CRuMs, the Opimoda, strongly supported in the LG+C60+G4 ML tree (99% npbs). However, in the BI analyses, Ancyromonadida appeared as the earliest-branching lineage within Opimoda (0.99 PP and 0.75 PP under CAT-GTR and CAT-Poisson models, respectively), Malawimonadida being sister to the Podiata clade (0.98 PP and 0.73 PP under CAT-GTR and CAT-Poisson models respectively) (Figure 1; Figure S3D-E).

To test whether the aforementioned uncertainties in our phylogenomic tree, i.e. the position of *Manchomonas* within Apusomonadida, *Fabomonas* within Ancyromonadida and that of Malawimonadida, could result from differences in evolutionary rates, we monitored the statistical support (ufbs) of the different alternative topologies for the specific incongruent splits (Figure 2A-C) after progressive removal of the fastest-evolving alignment sites (5% at a time). The monophyly of *Manchomonas* and *Apusomonas* was strongly supported until only 10-15% of the data remained and that of *Planomonas* and *Fabomonas* was virtually fully supported until less than 25% of sites remained (Figure 2D). In the case of malawimonads, we monitored the statistical support for the two observed topologies in our analyses, as well as the monophyly of malawimonads and, respectively, Discoba and Metamonada. We also tested for the monophyly of ancyromonads and Metamonada (Figure 2C). We followed, as control, the monophyly of Opisthokonta, fully supported until 10% of sites remained (Figure 2D). We recovered low statistical support for the two observed alternatives, although the monophyly of malawimonads+ancyromonads was always better supported than malawimonads+Podiata. By contrast, the monophyly of malawimonads and Discoba was never supported (Figure 2D). Approximately unbiased (AU) tests only rejected the malawimonad+discoban monophyly (Table S6). The likelihood of malawimonads being sister to metamonads or podiates was not significantly different from that of the malawimonads+ancyromonads monophyly in the best supported tree (Figure 1). Similarly, the monophyly of ancyromonads and metamonads cannot be discarded (Table S6). However, these two metamonad topologies were never supported by our fast-evolving removal analysis (Figure 2D). The inclusion of a more thorough taxonomic sampling branching deeply around these pivotal nodes should help solving problematic relationships, notably that of metamonads, and strengthen the support of deep nodes in the eukaryotic tree.

**Figure 2.**
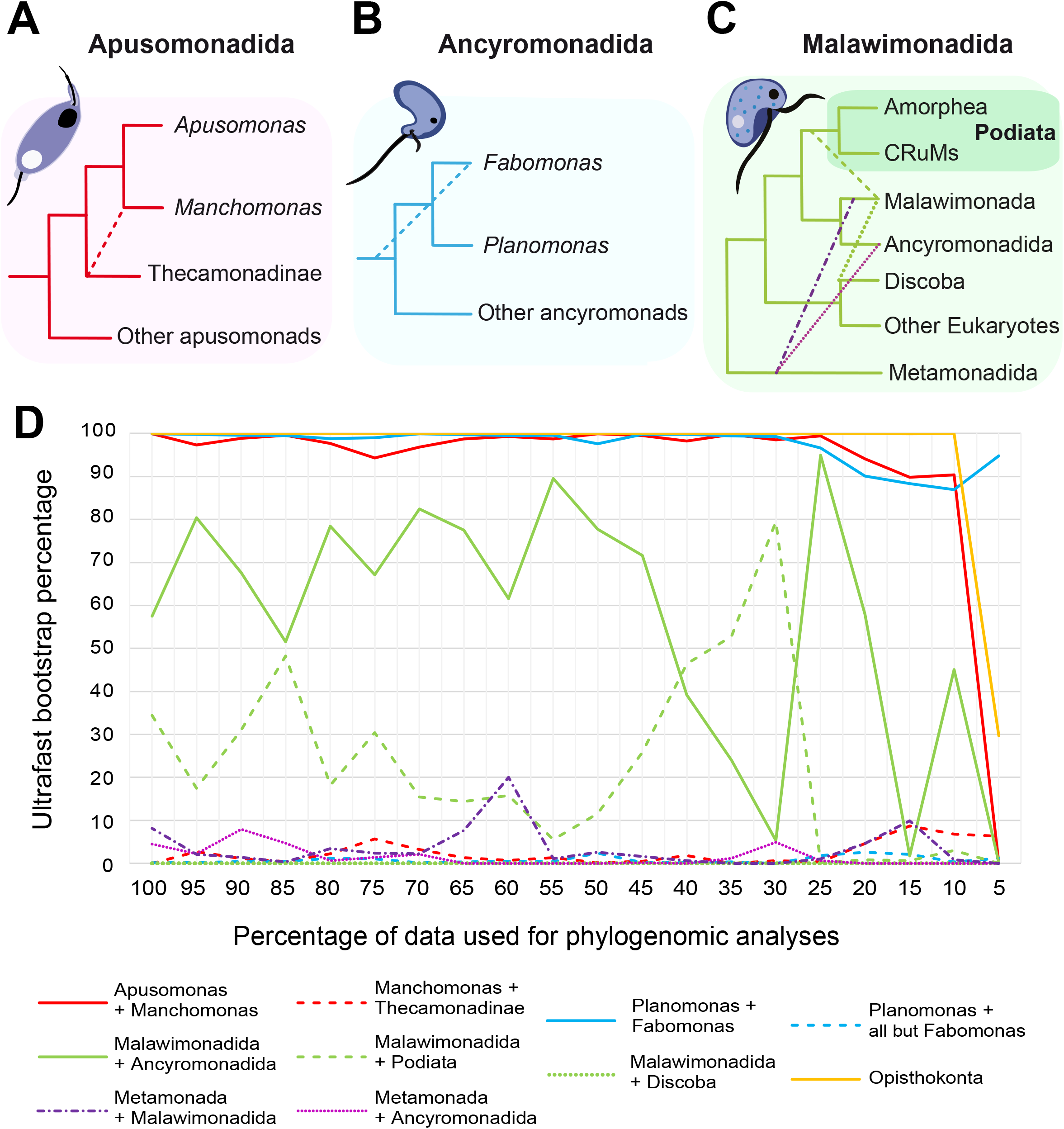
Progressive removal of the fastest evolving sites in 5% increments to evaluate alternative topologies within Opimoda. The alternative positions tested were as follows. **a**, Within the Apusomonadida: *Manchomonas* being sister to *Apusomonas* or to the Thecamonadinae (dashed line). **b**, Within the Ancyromonadida: the monophyly of *Fabomonas* and *Planomonas* (the Planomonadidae clade) versus their paraphyly (*Fabomonas* branching deeply, dashed line). **c**, Regarding Malawimonadida and Ancyromonadida: Malawimonadida and Ancyromonadida being monophyletic versus (dashed lines) Malawimonadida branching as sister to i) Podiata (and, accordingly, ancyromonads as the earliest-branching Opimoda lineage), ii) Discoba or iii) Metamonada; we also tested the monophyly of Ancyromonadida and Metamonada. **d**, Plot showing the bootstrap support in ML phylogenetic trees under the LG+C60+G4 model of sequence evolution as sites are progressively removed. The monophyly of Opisthokonta (yellow) is used as a control to indicate when the phylogenetic signal is too low to retrieve well-known robust monophyletic clades.

Although our trees were not rooted with an external outgroup, we included the widest possible diversity of free-living protists also on the Diphoda^19^ side, including heterotrophic flagellates when possible. We arbitrarily rooted our trees using the excavate clade Metamonada, including the relatively short-branching *Trimastix* and *Paratrimastix* species (Figure 1). As expected, Metamonada and Discoba appeared paraphyletic to each other^40,41^. Also, we recovered the monophyly of Diaphoretickes with overall high support (Figure 1). Within Diaphoretickes, the SAR supergroup also received full support. However, the monophyly of SAR and telonemids found in some analyses (the TSAR group^42^) was not observed. Telonemids branched within or sister to Haptista, albeit with moderate-to-low support (Figure 1). Archaeplastida were recovered with moderate (96%; ML) to full support (1 PP; BI CAT-GTR) and included Picozoa (with only BI CAT-GTR-based full support) as sister to Rhodelphida+Rhodophyta, as recently observed^43^. However, Picozoa appeared related to telonemids and haptophytes in ML trees with ELM PMSF (76%-npbs) and BI CAT-Poisson (0.5 PP; Figure S3C and E). We also recovered with full support the widely accepted monophyly of Archaeplastida and Cryptista^13,43^ using BI CAT-GTR. Finally, we retrieved the monophyly of *Ancoracysta* (Provora)^16^, Hemimastigophora^13^ and *Meteora sporadica* CRO19MET^17^ with full ML npbs support (Figure 1), as recently observed for two other *M. sporadica* strains^18^. *Ancoracysta* and other Provora members have also been suggested to form a monophyletic group with Hemimastigophora^15^. Our results thus provide additional support for this new supergroup of morphologically diverse predatory protists.

### Evolution of major phenotypic traits during the early opimodan radiation

This phylogenetic framework (Figure 1) allows to comparatively assess the distribution of complex phenotypic features, infer the ancestral states for the different clades and propose plausible evolutionary scenarios for trait evolution. Obviously, the availability of morphological and structural data is still limited, many morphological features may not be homologous, and more biodiversity alongside ultrastructural and phylogenetic analyses with, notably, the inference of a rooted tree of eukaryotes (e.g. using mitochondrial or Asgard archaeal-derived markers) will be needed to validate and/or complete this emerging evolutionary scheme (Figure 3).

**Figure 3.**
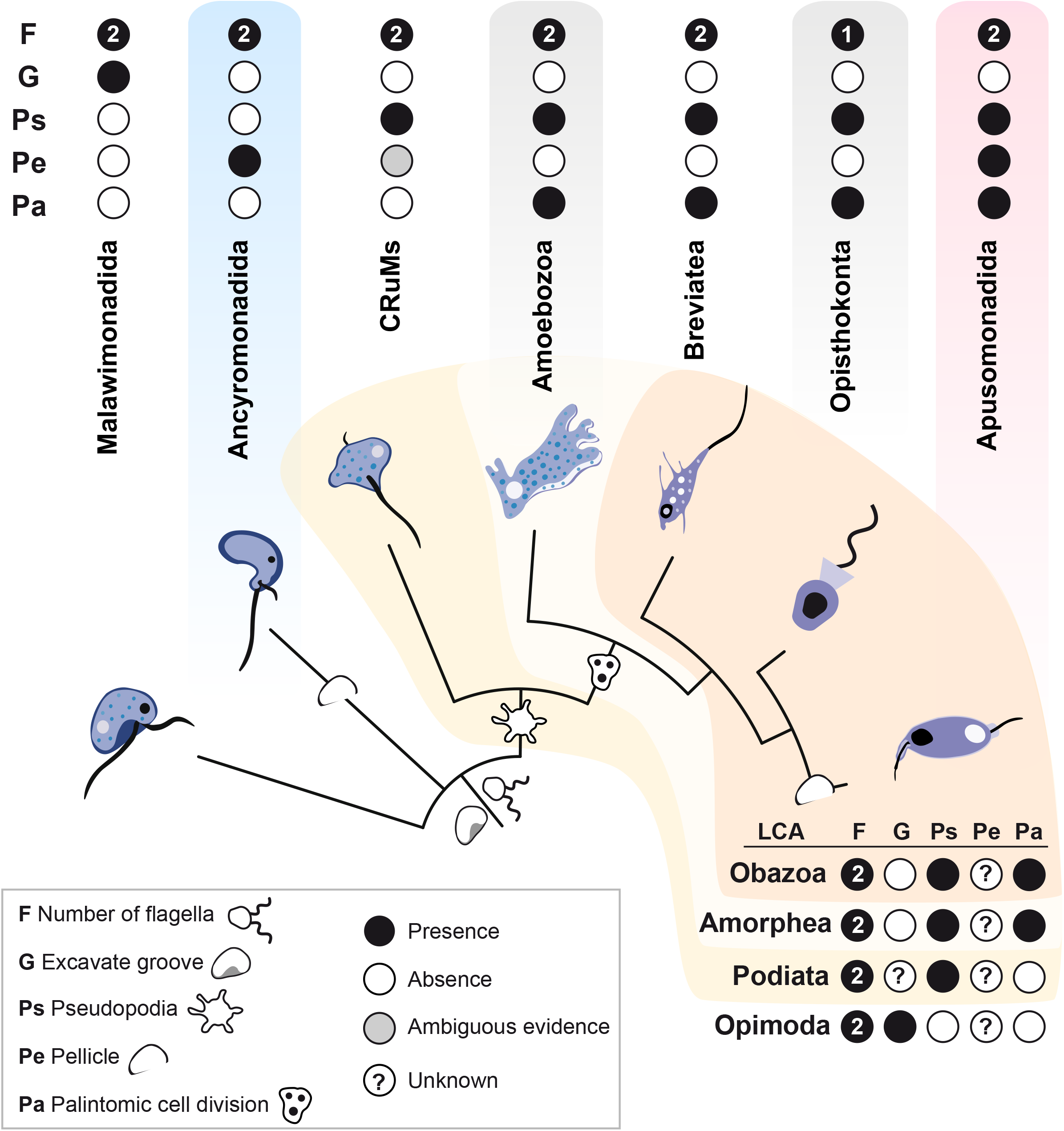
Early trait evolution across Opimoda lineages. The distribution of five key morphological traits is parsimoniously inferred for each lineage ancestor (upper panel) based on available descriptions and, to their respective last common ancestors (LCA), based on the inferred phylogenetic backbone (lower panel). Numbers in black circles refer to the number of flagella. The small character drawings in the cladogram represent the innovations at the onset of each lineage.

Interestingly, some large clades, notably the ancyromonads, show apparent phenotypic stasis, without obvious differences in cell size, shape or other observable feature under light microscopy^27^. With a genetic divergence comparable to other morphologically and structurally diverse lineages such as Amoebozoa or Opisthokonta, ancyromonads retained a constrained morphotype for hundreds of million years, suggesting an efficient adaptation to their predatory lifestyle in benthic and soil ecosystems. Ecologically, ancyromonads comprise marine and freshwater species, which belong to distinct clades for the current taxon sampling, suggesting a single transition from marine to freshwater ancestors (Figure 1). Accordingly, the last ancyromonad common ancestor was likely marine and resembled extant species, having bean-shaped flattened biflagellate cells with a short anterior flagellum and a rostrum with extrusomes^27^. Although ancyromonads possess a dorsal pellicle, a complex cytoskeleton and a ventral groove for feeding, they lack the bona fide excavate groove or actin-based pseudopods similar to those of diverse Obazoa^27^ (Figure 3). By contrast, apusomonads exhibit some morphological differences alongside ecological variation, with marine-freshwater transitions having occurred multiple times during their evolution^24,26^ (Figure 1). Clade A members exhibit a short proboscis sleeve compared to cell body length, and more-prominent pseudopodia than other apusomonads^26^. *Podomonas* has a tusk, a character also present in Thecamonadinae^44^. *Multimonas* displays a potential tusk, posterior extrusomes^45^ and division by binary and multiple fission. *Podomonas* and *Mylnikovia* possess refractile granules running in parallel to the posterior flagellum^46^. All these characteristics can be found in other apusomonads^26^. In clade B, if *Karpovia* or *Singekia* present a proper tusk (still unclear) alongside trailing pseudopodia^26^, they would fit the subfamily Thecamonadinae description^44^. *Manchomonas* and *Apusomonas* share the presence of few pseudopods, a hidden posterior flagellum, the absence of tusks, and some ultrastructural features^47^. *Manchomonas* has the largest observed sleeve compared to other elongated apusomonads, while *Apusomonas* exhibits a unique structure called mastigophore. The elongated, *Amastigomonas*-type, cell shape seems the ancestral apusomonad phenotype, the rounded *Apusomonas* phenotypes being derived. The last apusomonad common ancestor likely had elongated cell type, dorsal pellicle, ventral feeding groove, actin-based pseudopodia, proboscis and possibly a tusk, and was able to divide by multiple fission (Figure 3).

In our phylogenomic tree, malawimonads and ancyromonads were sister groups (Figure 1). If confirmed, this clade would be one of the earliest-branching lineages after the Opimoda-Diphoda split. Alternatively, malawimonads could be sister to the Podiata. Whatever the actual topology, since both clades encompass small bacterivorous heterotrophic biflagellates with a, likely homologous^48^, ventral feeding groove^12,27,49^, the last common ancestor of Opimoda most likely shared this excavate-like phenotype (Figure 3). Furthermore, since excavates are likely paraphyletic^14,15,50^, with Metamonada and Discoba branching deeply in the eukaryotic tree (Figure 1), the ancestral LECA likely shared that excavate-like biflagellate phenotype^51,52^, irrespective of the specific position of the root^53^ (Figure 3). An excavate-like ancestor seems further supported by the strong conservation of the microtubular cytoskeleton and the flagellar apparatus^54^ across eukaryotes^55^. From such an ancestor, ancyromonads developed a dorsal pellicle and a rostrum with extrusomes, losing the flagellar vanes and shortening the anterior flagellum^12^. The opimodan ancestor likely was a marine planktonic biflagellate that adapted to glide on benthic substrates and, in subsequent evolution, lost free-swimming capabilities. Without the excavate physical constrains, feeding being intertwined with the flagellar motility (the excavate groove hosting the posteriorly-oriented flagellum)^48^, and without highly developed actin-based pseudopodia, the ancyromonad morphotype might have easily evolved (Figure 3). Bearing a dorsal pellicle, ancyromonads maintained ventral feeding, keeping the posterior flagellum attached to the surface, which is likely at the origin of their “twitch-yanking” feeding movement^27^.

Our phylogenomic tree supported the monophyly of Amorphea (Amoebozoa, Breviatea, Apusomonadida and Opisthokonta) and CRuMs (Figure 1). This clade, named Podiata^34^, had already been proposed on the shared capability to produce pseudopodia. The last Amorphea common ancestor was clearly able to produce pseudopodia (Figure 3). However, cell biology descriptions for CRuMs are still limited^14^. Accordingly, the presence of an apusomonad/ancyromonad-homologous pellicle^56^, or the presence of true pseudopodia (e.g., in *Micronuclearia* and *Rigifila*), needs confirmation. Nonetheless, some *Mantamonas* species (e.g. *M. plastica*) bear pseudopodia^57^ and, therefore, the ancestor of the group likely did so. Mantamonadida have a stable phylogenetic position^58^ and might have retained some of the ancestral features of CRuMs; indeed, *Mantamonas* share some similarities with ancyromonads, notably the possession of small flattened cells with a short anterior flagellum and benthic/soil-associated lifestyles. However, the other CRuM linages (Diphyllatea and Hilomonadea) include larger, freshwater planktonic species, also possessing a wider, non-excavate, ventral groove^56,59^. Populating the CRuMs’ branch with new described members should help to ascertain trait evolution within this clade.

The Amorphea contain the most diversified and studied lineages of the opimodan side of the eukaryotic tree, Amoebozoa and Opisthokonta. These clades exhibit more diverse and derived morphoplans than the two other amorphean lineages, Apusomonadida and Breviatea. The amoebozoan ancestor was biflagellated and possessed a complex flagellar apparatus^60^. However, many Amoebozoa and the Opisthokonta lost one or both flagella^61,62^. Biflagellate protists have complex microtubular cytoskeletons that impose severe structural constrains on their cell shape, such that flagellar loss, freeing those constrains, likely facilitated the evolution of more diverse morphoplans. This increased morphological evolvability linked to concomitant changes in selective pressures allowed the exploration of novel cell shapes (e.g. amoeboid), feeding modes (e.g., osmotrophy), and cell-cell interactions (e.g., multicellularity), as currently observed across these clades^39^. By contrast, Breviatea and Apusomonadida, paraphyletic within the Obazoa, likely retained the ancestral biflagellate state alongside other features likely present in the obazoan ancestor (Figure 3). Both lineages encompass small bacterivorous amoeboflagellates that phagocytize prey using pseudopodia (unlike ancyromonads or excavates)^46^. Breviates are anaerobic and possess gliding and swimming forms, lack a pellicle and display more pronounced amoeboid shapes than apusomonads^22^. Apusomonads comprise gliding, elongated, semi-rigid amoeboflagellates with an anterior proboscis and a dorsal pellicle allowing ventral feeding similarly to ancyromonads^26,44^. Interestingly, both breviates and apusomonads can divide palintomically, i.e. their karyokinesis is uncoupled from cellular division, leading, at least transiently, to multinucleated cells. The ability to generate multinucleated cells is also widespread across amoebozoans and opisthokonts, including several unicellular relatives of animals (e.g. corallochytrids)^63^ and fungi (e.g. aphelids)^20^. It seems increasingly clear that the occurrence of coenocytic (i.e., multinucleated) stages was important along the road to metazoan multicellularity^21^. Therefore, the ancestral Amorphea capability to produce multinucleated cells appears intriguingly crucial for the subsequent evolution of plasmodial growth, and metazoan and hyphal-based fungal multicellularity.

## STAR★Methods

### Key resources table

### Resource availability Lead contact

Further information and requests for resources and reagents should be directed to and will be fulfilled by the Lead Contact, Purificación López García.

### Materials availability

This study did not generate new unique reagents.

### Data and code availability

Raw read sequences have been submitted to Sequence Read Archive under BioProject PRJNA907040; the specific accession numbers for each protist transcriptome are given in Table S1. Transcripts and oligopeptides generated in this study are available in the figshare repository (10.6084/m9.figshare.22148027).

## Method details

### Protist culture, RNA extraction and transcriptome sequencing

The 22 species of heterotrophic flagellate protists from which we generated transcriptome data were previously isolated or enriched from benthos of marine or freshwater ecosystems or soil, and included apusomonads^26^, ancyromonads^27^ and *Meteora sporadica strain* CRO19MET^17^, cultured from Mediterranean benthic samples in the same area as the original type strain^30^ (Table S1; Figure S1). Soil and freshwater flagellates were grown in Volvic water with 1% yeast tryptone (YT), and marine flagellates in 0.2 micron-filtered seawater with 1% YT medium or Cerophyl medium. Flagellates were grown in flat cell culture flasks with ∼10 ml of medium, just to cover the bottom surface (75 cm^2^). The gliding protist cells from high-density cultures were collected by gently scratching the bottom of the flask with a cell scraper and pooled in 50 ml Falcon tubes. Cells were pelleted by centrifugation at 10ºC for 15 minutes at 15,000g. Total RNA for each species was extracted with the RNeasy mini Kit (Qiagen) following the manufacturer protocol, including DNAse treatment. Purified RNA was quantified using a Qubit fluorometer (ThermoFisher Scientific). For each species, cDNA Illumina libraries were constructed after polyA mRNA selection, tagged and paired-end (2 × 150 bp) sequenced with Illumina NovaSeq 6000 S2 (Eurofins Genomics, Germany) in three different sequencing runs (NG-22350, NG-24277, NG-25209; Figure S2 and Table S3). Sequence statistics and accession numbers are provided in Table S1.

### Transcriptome assembly and decontamination

Our bioinformatic pipeline is summarized in Figure S2. The quality of Illumina sequences was checked with FastQC^64^ v0.11.8. High-quality reads were retained and used for transcriptome *de novo* assembly using Spades^65^ v3.13.1 with -rna mode and default parameters. The raw transcripts were used for phylogenetic marker selection (see next section). Cross-contamination of transcripts among multiplexed cDNA libraries within the same sequencing run was detected and removed using CroCo^66^ v1.2 (Figure Figure S2, Tables S2-S3). We translated the remaining transcripts into oligopeptides using TransDecoder v5.5 (https://github.com/TransDecoder/) with the script LongOrfs and clustered them with CD-HIT^67^ v4.8.1 at 100% identity. Predicted oligopeptides of non-eukaryotic origin were removed using similarity search against a custom protein database (BAUVdb: bacteria, archaea, eUkaryotes and viruses). BAUVdb included 32 reference genomes of eukaryotic species spanning all supergroups (Table S4) as well as, respectively, 62,291 and 3,412 bacterial and archaeal genomes from the Genome Taxonomy DataBase, GTDB^68^ release 207, 361,930 viral genomes from the reference viral database (RVDB)^69^ and RVDB proteins^70^ clustered using CD-HIT at 90% identity and coverage. To discriminate target eukaryotic sequences from those of alien origin, we then applied Diamond^71^ v2.0.14.152 in ultra-sensitive mode setting an e-value threshold of 1e-3 for a maximum of 100 hits. From the tabular output, eukaryotic oligopeptides were retained if *Thecamonas trahens* or five eukaryotes were the first hits. Eukaryotic oligopeptides were automatically annotated using eggNOG-mapper^72^ using all orthologs and all annotations as evidence. The number of identified single-copy markers from BUSCO^31^ v5.2.2 was used as a proxy for gene set completeness using the eukaryote database db10. All transcripts and oligopeptides generated in this study are available in the figshare repository (10.6084/m9.figshare.22148027). Since *A. proboscidea* MPSANABRIA15 was co-cultured with a difficult-to-eliminate stramenopile contaminant (belonging to the genus *Paraphysomonas*, as inferred from its 18S rRNA gene sharing ∼98% pairwise identity with members of this genus), we also sequenced the transcriptome of the latter from a monoeukaryotic culture (NCBI biosample SRR23610779). To decontaminate the apusomonad set of proteins from stramenopile sequences, eukaryotic proteins of *A. proboscidea* MPSANABRIA15 were used as queries in BLASTP (minimum e-value of 1e-25) against a protein database with the closest related taxa (PROMEX), our own stramenopile contaminant, plus three publicly available *Paraphysomonas* transcriptomes: MMETSP1103, MMETSP0103 and MMETSP1107. Eukaryotic oligopeptides were considered contaminant when they had only stramenopile hits or, in the case of hits in both lineages, when the pairwise identity, weighted by coverage, was higher in stramenopiles than in PROMEX.

### Phylogenomic analyses

Our phylogenomic dataset was updated from a previous paneukaryotic study with 104 taxa and 351 conserved markers^13^. To that dataset, we added the corresponding identified markers from recently sequenced genomes/transcriptomes of early-branching eukaryotes^42,73-76^ and from our 22 flagellate raw transcriptomes translated into oligopeptides using TransDecoder, i.e. without undergoing any cleaning step (Table S5). To identify the selected phylogenetic markers in these transcriptomes, we queried these datasets with the 351 marker sequences from *Homo, Saprolegnia, Spizellomyces* and *Diphylleia* with BLASTp and retrieved all possible homologs. Each marker from the 230 taxa representing all eukaryotic supergroups, was aligned with MAFFT^77^ v7.427 (L-INS-i with 1,000 iterations), and trimmed using Trimal^78^ v1.4.rev22 in automated mode. Approximate-maximum likelihood single marker trees were inferred from each trimmed alignment using FastTree^79^ v2.1.11 with default parameters, and examined manually with Figtree v1.4.3 (http://tree.bio.ed.ac.uk/software/figtree/). Alignments were inspected with Aliview^80^ v1.26 80, and pruned from contaminants, cross-contaminants, fragmented sequences, paralogs or spuriously aligned sequences. A total of 48 markers with complex histories were removed from the dataset, resulting in a selection of 303 protein-coding genes. For final multi-marker phylogenetic analyses and to limit computational load, the taxon sampling was reduced to 101 eukaryotes, keeping at least five representatives for each known supergroup, mostly free-living heterotrophic flagellates (available in the figshare repository; 10.6084/m9.figshare.22148027). Finally, the markers were realigned with MAFFT L-INS-I with 1,000 iterations, trimmed with Trimal in automated mode and concatenated with alvert.py from the barrel-o-monkeys (http://rogerlab.biochemistryandmolecularbiology.dal.ca/Software/Software.htm#Monkeybarrel), generating a concatenated matrix with 97,171 amino acid sites that was used to infer phylogenies (10.6084/m9.figshare.22148027). The Maximum likelihood (ML) phylogenetic tree was inferred using IQ-TREE^81^ v1.6. We first generated a guide tree inferred with the LG+C60+G4 mixture model (-mwopt flag), and then generated a PMSF model^82^. The statistical support was generated with 1,000 ultrafast bootstraps without PMSF (Figure S3A) and with 100 non-parametric bootstrap under the PMSF model (Figure 1). We also reconstructed an ML tree using the recently implemented, eukaryote-adapted ELM model^32^ with 1,000 ufbs without PMSF (Figure S3B), and with 100 npbs under the PMSF model (Figure S3C). The Bayesian inference (BI) analyses were conducted with PhyloBayes-MPI^83^ v1.5a with both CAT-GTR (Figure S3D) and CAT-Poisson (Figure S3E) models^84^, with two MCMC chains respectively run for 10,000 generations, saving one every 10 trees (although four MCMC chains were run for the CAT-GTR model, only two properly converged, as shown using Tracer^85^; Figure S4). Analyses were stopped once convergence thresholds were reached (i.e. maximum discrepancy <0.1 and minimum effective size >100, calculated using bpcomp) and consensus trees constructed after a burn-in of 25%. Additionally, to minimize possible systematic errors, the fastest-evolving sites were progressively removed at 5% of sites at a time. For that, among-site substitution rates were inferred using IQ-TREE under the -wsr option and the best-fitting model, generating a total of 19 new data subsets (10.6084/m9.figshare.22148027). Each of them was used to infer a phylogeny using IQ-TREE with the LG+C60+G4 model and obtain the ufbs for each split. We used CONSENSE from the PHYLIP v3.695 package (https://phylipweb.github.io/phylip/) to interrogate the ufboot files using a Python script (Nick Irwin, pers. comm.) and recover the support for each specified split (Figure 2). Approximately unbiased (AU) tests were conducted as explained in the IQ-TREE documentation. We first used Mesquite^86^ to create star trees with a constrained split for the topology to be tested before inferring ML trees with IQ-TREE with the –z and –au options (Tables S6).

## Acknowledgements

We are grateful to Tom Cavalier-Smith and Emma Chao for their contribution to evolutionary protistology and their friendly availability to discuss ideas and share protist strains. We thank Ares Rocañín-Arjó, Saioa Manzano, Moisès Bernabeu, Xavier Grau-Bové and Nick Irwin for bioinformatic insights. We thank the UNICELL platform (http://www.deemteam.fr/en/unicell) for help in transcriptome production. This work was supported by the European Research Council (ERC) Advanced Grants ‘Protistworld’ and ‘Plast-Evol’ (322669 and 787904, respectively) and the Horizon 2020 research and innovation programme under the Marie Skłodowska-Curie ITN project SINGEK (http://www.singek.eu/; grant agreement no. H2020-MSCA-ITN-2015-675752). G.T. was supported by 2019 BP 00208, Beatriu de Pinós-3 Postdoctoral Programme (BP3), Grant agreement ID: 801370. LG was funded by the Horizon 2020 research and innovation programme under the European Marie Skłodowska-Curie Individual Fellowship H2020-MSCA-IF-2020 (grant agreement no. 101022101 ? FungEye).

