## Supplementary Information for "Phylogenomics of neglected flagellated protists supports a revised eukaryotic tree of life"

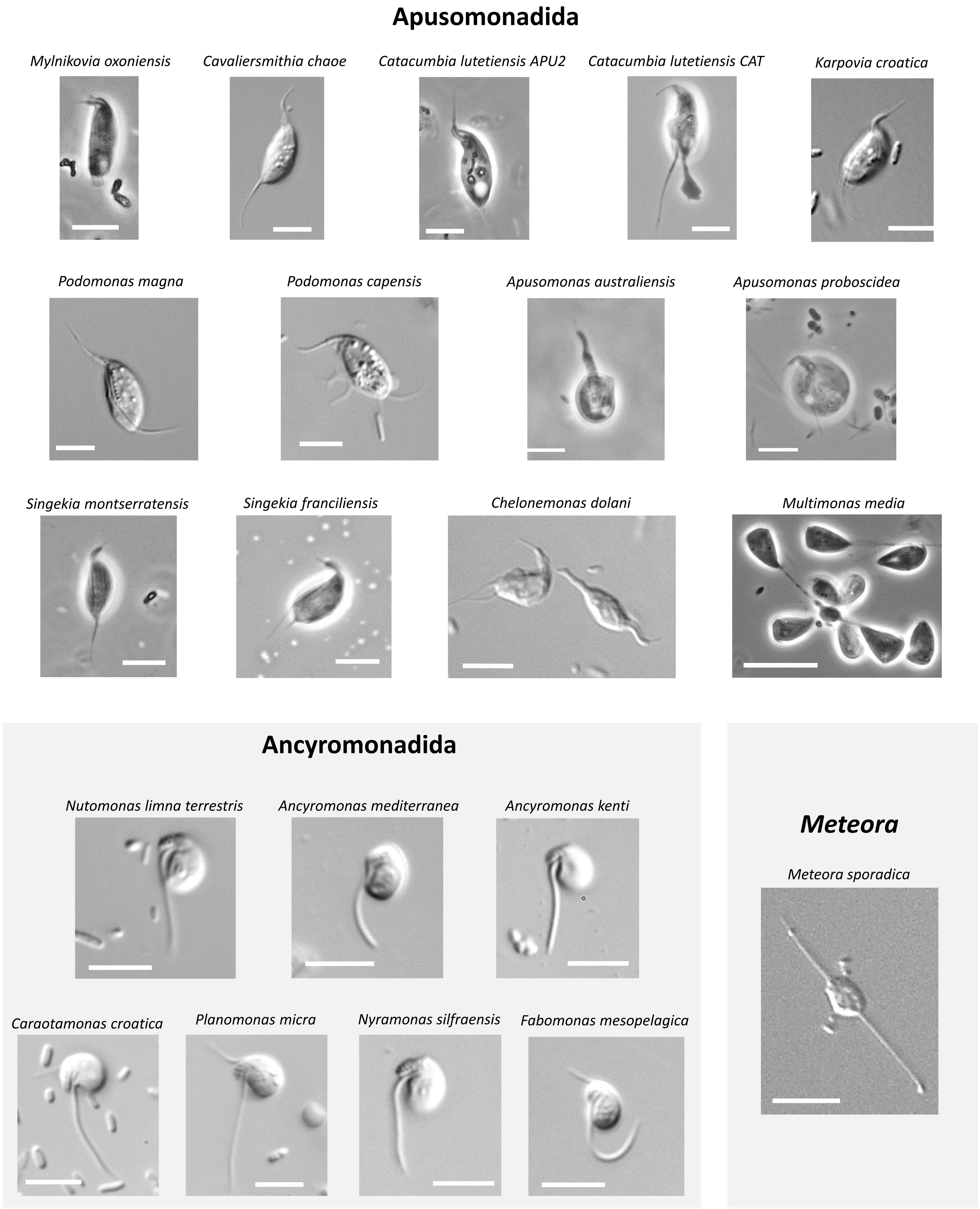


**Figure S1. Optical microscopy photographs of the protists cultured from diverse marine and freshwater benthic samples from which transcriptomes were generated.** The strain identifiers are provided in Table S1. A full description of the apusomonad, ancyromonad an *Meteora* strains cultured from environmental samples in our lab is given in, respectively, refs. 26, 27 and 17. Images and data about Chelonemonas geobuk can be found in ref. 25. The scale bar corresponds to 5 µm.

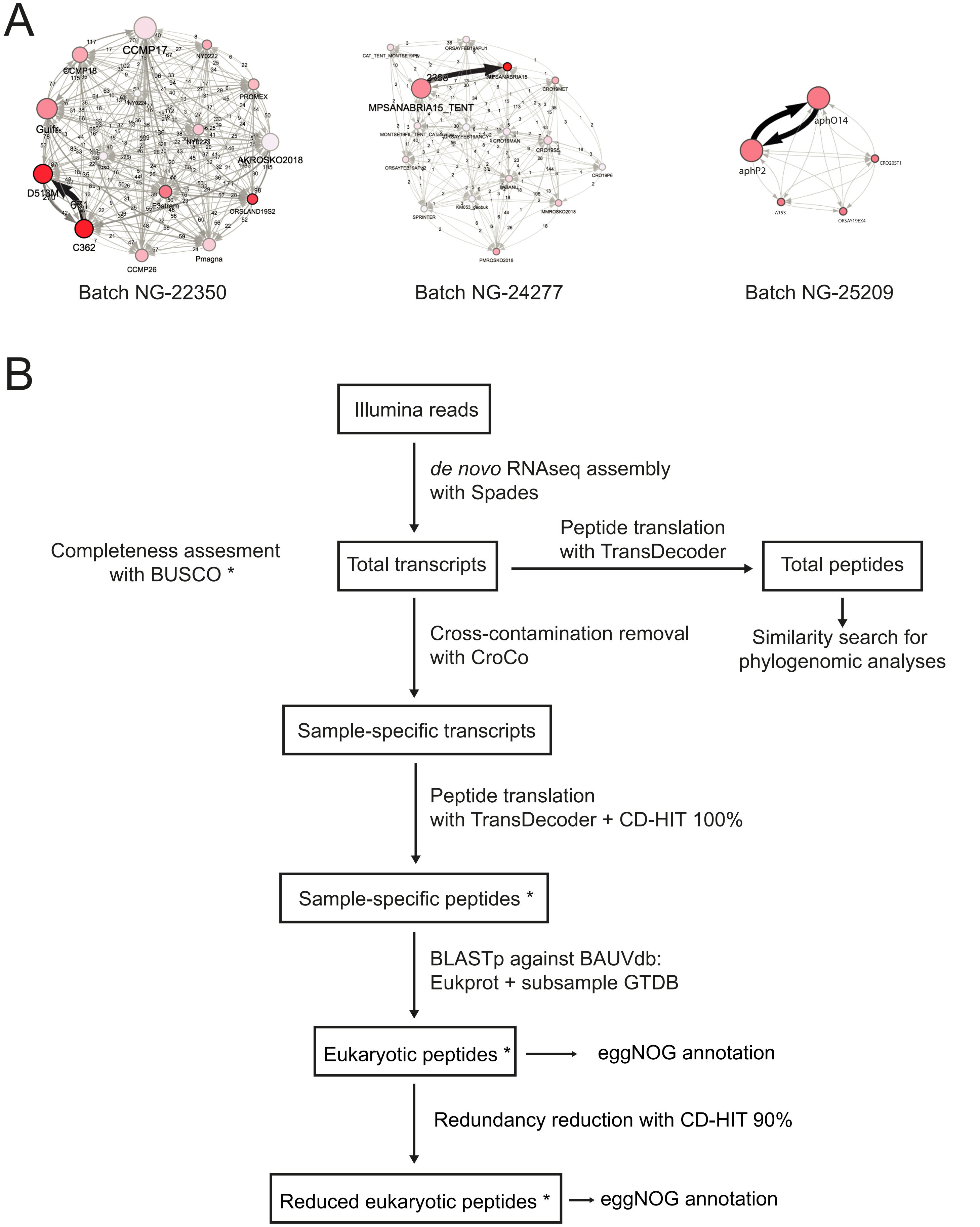


**Figure S2. RNAseq decontamination process. A.** Cross-contamination networks within batch samples NG-11350, NG-24277 and NG-25209 (see Table S3). **B.** Bioinformatic pipeline to clean RNA sequences (see Tables S2 and S4).

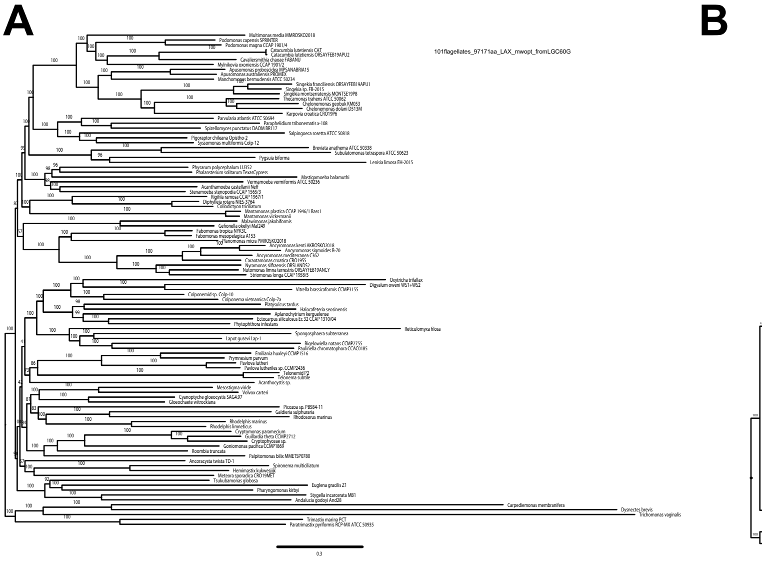

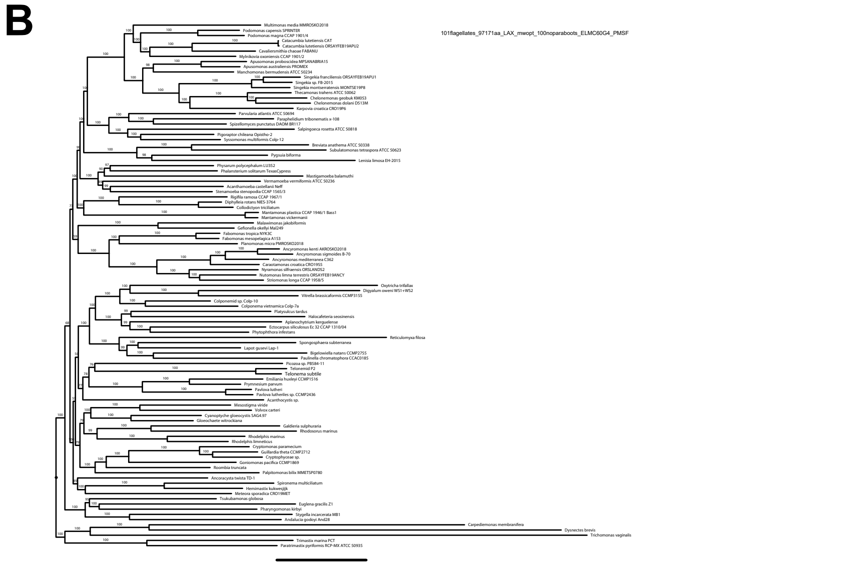

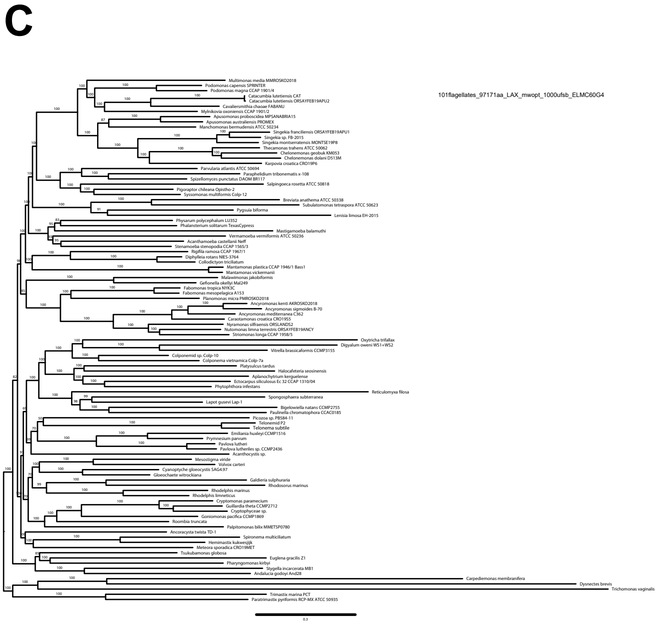

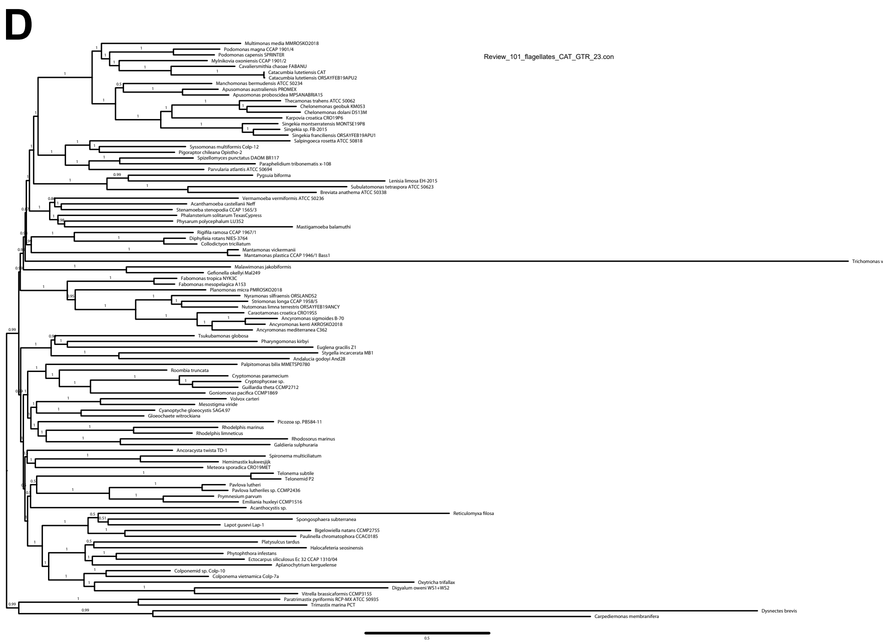

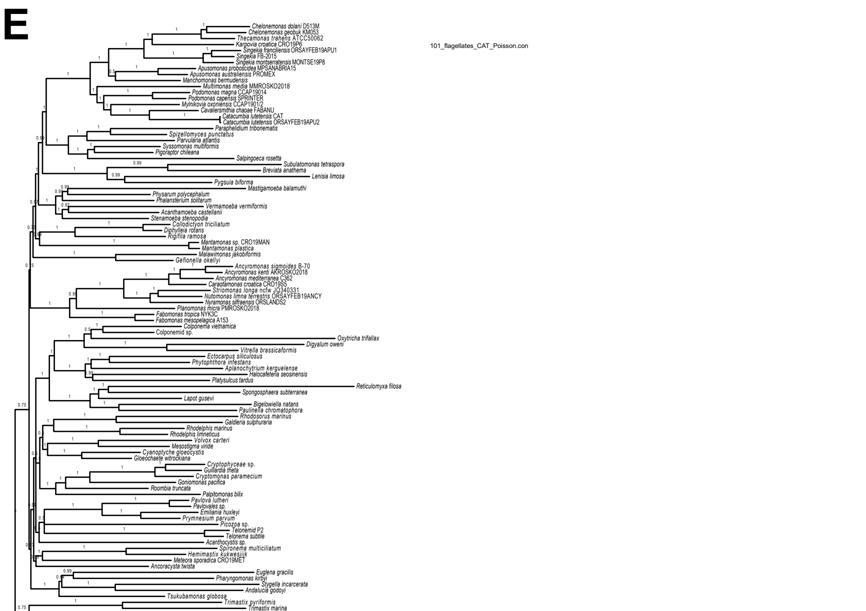

**Figure S3.** Phylogenomic trees of eukaryotes enriched in deep-branching protists. The trees were reconstructed using 303 conserved proteins and 97,171 conserved amino acid positions for a total of 101 species. **A.** Tree inferred with IQ-TREE using the model LG+C60+G4 with 1,000 ultrafast bootstraps as statistical support. **B.** Tree inferred with IQ-TREE using the PMSF approximation of the model ELM+C60+G4 with 100 non-parametric bootstraps as statistical support. **C.** Tree inferred with IQ-TREE using the model ELM+C60+G4 with ultrafast bootstraps as statistical support. **D.** Tree inferred with PhyloBayes under the CAT-GTR model, consensus of two chains, with posterior probabilities as statistical support. **E.** Tree inferred with PhyloBayes under the CAT-Poisson model, consensus of two chains, with posterior probabilities as statistical support.

**A**

| Summary Statistic | chain_1.tracetime | chain_2.tracetime | chain_3.tracetime | chain_4.tracetime |
|---|---|---|---|---|
| mean | 1838.8424 | 1778.6912 | 1879.5296 | 1966.3407 |
| stderr of mean | 172.3711 | 153.0129 | 217.5938 | 67.9229 |
| stdev | 567.5938 | 534.4329 | 596.7908 | 398.7175 |
| variance | 3.2216E5 | 2.8562E5 | 3.5616E5 | 1.5898E5 |
| median | 1965.912 | 1866.19 | 2001.674 | 1976.475 |
| value range | [701.349, 3758.… | [709.435, 1399… | [754.33, 6761.0… | [780.623, 4146.… |
| geometric mean | 1735.0566 | 1686.0522 | 1763.3296 | 1922.0836 |
| 95% HPD interval | [738.186, 2578.… | [713.735, 2448.… | [778.476, 2645.… | [816.905, 2519.… |
| auto-correlation time (ACT) | 876.7007 | 776.8559 | 1241.3736 | 270.353 |
| effective sample size (ESS) | 10.8 | 12.2 | 7.5 | 34.5 |
| number of samples | 9505 | 9476 | 9337 | 9315 |

**B**

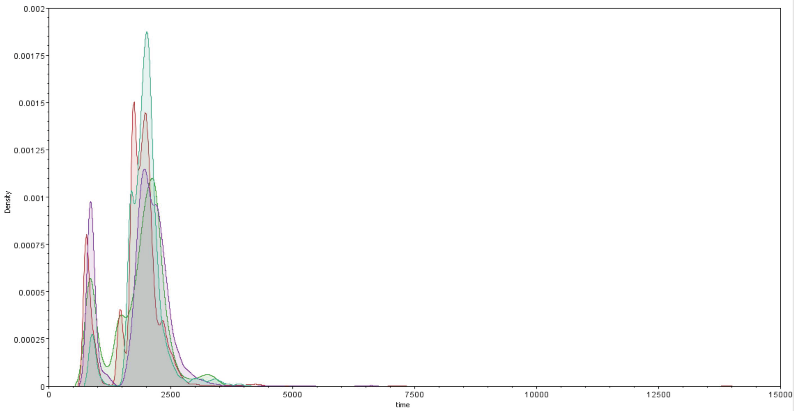


**C**

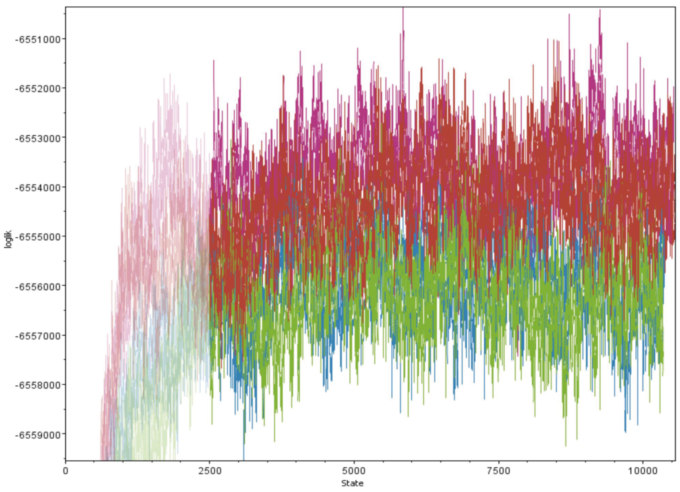


**D**

| Summary Statistic | chain_1.tracetime | chain_2.tracetime |
|---|---|---|
| mean | 914.1476 | 991.3041 |
| stderr of mean | 44.9764 | 43.2141 |
| stdev | 172.9599 | 170.0396 |
| variance | 29915.1347 | 28913.4503 |
| median | 865.249 | 944.52 |
| value range | [597.108, 1974.… | [741.757, 1942.… |
| geometric mean | 900.2334 | 978.7438 |
| 95% HPD interval | [663.876, 1271.… | [800.631, 1358.… |
| auto-correlation time (ACT) | 899.2165 | 871.1631 |
| effective sample size (ESS) | 14.8 | 15.5 |
| number of samples | 13297 | 13487 |

**E**

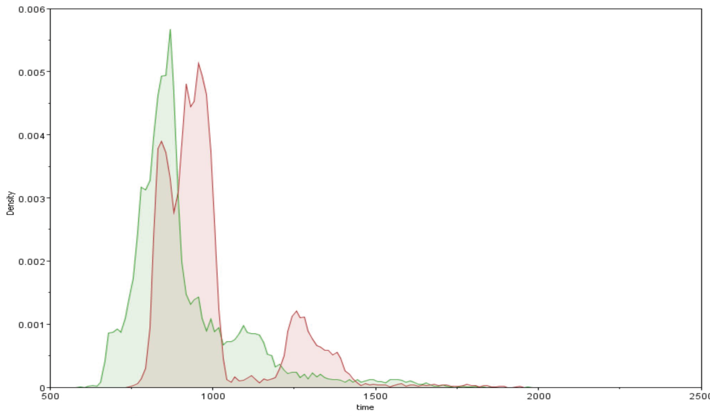


**F**

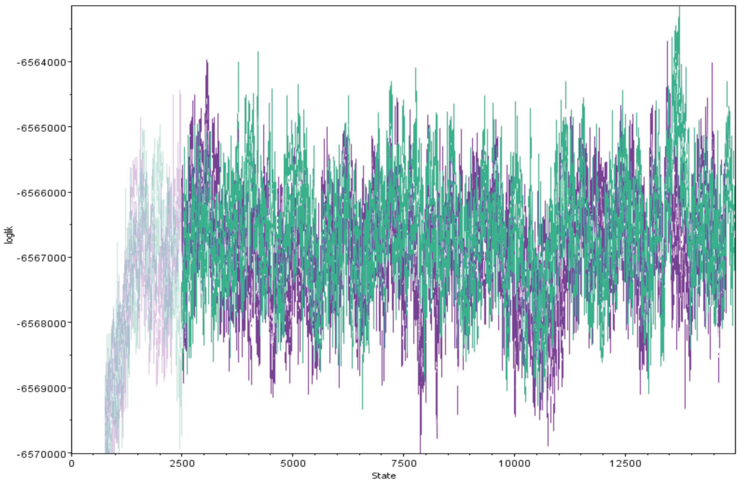


**Figure S4. Tracer analysis of Bayesian MCMC chains.** Screenshots of the Tracer analysis of the four MCMC chains from the PhyloBayes CAT-GTR run (A - C), and two MCMC chains from the PhyloBayes CAT-Poisson run (D - F). A and D show the statistical estimates; B and E, the marginal densities; C and F, the combined log likelihood per step.

**Table S1. Statistics of the transcriptomes sequences generated for several protist species.** The dataset includes 14 apusomonad and 7 ancyromonad species, as well as the *Meteora sporadica* (the type species of the genus *Meteora* ). PE, pair-ends. Data are provided for all eukaryotic transcripts and for transcripts clustered at 90% identity using cd-hit in order to eliminate paralogs/isoforms. Completion % refers to the percentage of BUSCO markers identified in the OrthoDB ortholog database.

| Species | Isolate/Strain | Environmental source | Accession number | No. of PE reads | All eukaryotic transcripts | | Non-redundant transcripts | |
|---|---|---|---|---|---|---|---|---|
| | | | | | No. of oligopeptides | Completion % BUSCO | No. of oligopeptides | Completion % BUSCO |
| Apusomonadida (14 spp.) | | | | | | | | |
| *Mylnikovia oxoniensis* | IVY8c CCAP1901/2 | Freshwater | SAMN32031840 | 33928521 | 19785 | 82.4 | 11455 | 82.4 |
| *Cavaliersmithia chaoae* | FABANU CCAP1346/1 | Marine | SAMN32031841 | 6183885 | 10130 | 90.6 | 9161 | 90.6 |
| *Catacumbia lutetiensis* APU2 | ORSAYFEB19APU2 | Freshwater | SAMN32031842 | 5236407 | 8090 | 89.4 | 7905 | 89.4 |
| *Catacumbia lutetiensis* CAT | CAT | Freshwater | SAMN32031843 | 5002746 | 7923 | 88.3 | 7713 | 88.3 |
| *Podomonas magna* | JJP-2003 CCAP1901/4 | Marine | SAMN32031844 | 19652204 | 12381 | 88.6 | 11559 | 88.6 |
| *Podomonas capensis* | SPRINTER | Marine | SAMN32031845 | 5028944 | 11266 | 89.1 | 10692 | 89.1 |
| *Multimonas media* | MMROSKO2018 | Marine | SAMN32031846 | 5365203 | 11967 | 84.6 | 9513 | 84.7 |
| *Apusomonas australiensis* | PROMEX | Freshwater | SAMN32031847 | 17927843 | 14430 | 90.6 | 12194 | 90.6 |
| *Apusomonas proboscidea* | MPSANABRIA15 | Freshwater | SAMN32031848 | 6170697 | 9192 | 63.9 | 9076 | 63.9 |
| *Karpovia croatica* | CRO19P6 | Marine | SAMN32031849 | 6682786 | 9291 | 88.2 | 9096 | 88.2 |
| *Singekia montserratensis* | MONTSE19P8 | Freshwater | SAMN32031850 | 5560582 | 9075 | 89.4 | 8943 | 89.4 |
| *Singekia franciliensis* | ORSAYFEB19APU1 | Freshwater | SAMN32031851 | 7738300 | 9101 | 89 | 8685 | 89 |
| *Chelonemonas dolani* | D513M | Marine | SAMN32031852 | 17648679 | 11352 | 92.9 | 11128 | 92.9 |
| *Chelonemonas geobuk* | KM053 | Marine | SAMN32031853 | 5642089 | 12141 | 92.9 | 11963 | 92.9 |
| Ancyromonadida (7 spp.) | | | | | | | | |
| *Nutomonas limna terrestris* | ORSAYFEB19ANCY | Freshwater | SAMN32031854 | 5921752 | 5514 | 55.7 | 5322 | 55.7 |
| *Caraotamonas croatica* | CRO19S5 | Marine | SAMN32031855 | 6157385 | 5552 | 80.4 | 5491 | 80.4 |
| *Ancyromonas mediterranea* | C362 | Marine | SAMN32031856 | 34510209 | 5807 | 80.9 | 5569 | 80.9 |
| *Ancyromonas kenti* | AKROSKO2018 | Marine | SAMN32031857 | 20255134 | 5583 | 78.1 | 5308 | 77.6 |
| *Planomonas micra* | PMROSKO2018 | Marine | SAMN32031858 | 6485357 | 6065 | 82 | 5953 | 82 |
| *Nyramonas silfraensis* | ORSLANDS2 | Freshwater | SAMN32031859 | 23609535 | 5764 | 81.2 | 5577 | 81.2 |
| *Fabomonas mesopelagica* | A153 | Marine | SAMN32031860 | 7208640 | 6662 | 89.1 | 6519 | 89.1 |
| Meteora | | | | | | | | |
| *Meteora sporadica* | CRO19MET | Marine | SAMN32031861 | 5936635 | 5197 | 87.5 | 4964 | 87.4 |

**Table S2.** Intermediate statistics during data processing for the protists transcriptomes generated in this study

| Group | Organism | Sp code | Isolate | Source | Accession | Batch | No. PE reads | BRUT: from *de novo* assembly – assembled transcripts: No. transcripts | BRUT: from *de novo* assembly – assembled transcripts: BUSCO | BRUT: from *de novo* assembly – predicted peptides + cdhit1: No. Oligopeptides | BRUT: from *de novo* assembly – predicted peptides + cdhit1: BUSCO |
|---|---|---|---|---|---|---|---|---|---|---|---|
| 14 Apusomonadida | *Mylnikovia oxoniensis* | MylOxo | IVY8c CCAP1901/2 | Freshwater | SAMN32031840 | NG-22350 | 33928521 | 78978 | 97.7 C:96.4%[S:32.0%,D:64.4%],F:1.3%,M:2.3% | 151623 | 98.1 C:96.5%[S:56.9%,D:39.6%],F:1.6%,M:1.9%,n:255 |
| 14 Apusomonadida | *Cavaliersmithia chaoae* | CavCha | FABANU CCAP1346/1 | Marine | SAMN32031841 | NG-24277 | 6183885 | 37052 | 97.7 C:94.4%[S:85.5%,D:8.9%],F:3.3%,M:2.3% | 64936 | 96.1 C:91.4%[S:90.6%,D:0.8%],F:4.7%,M:3.9%,n:255 |
| 14 Apusomonadida | *Catacumbia lutetiensis* APU2 | CatApu | ORSAYFEB19APU2 | Freshwater | SAMN32031842 | NG-24277 | 5236407 | 39106 | 95.7 C:90.1%[S:87.1%,D:3.0%],F:5.6%,M:4.3% | 57433 | 93.8 C:84.8%[S:82.4%,D:2.4%],F:9.0%,M:6.2%,n:255 |
| 14 Apusomonadida | *Catacumbia lutetiensis* CAT | CatLut | CAT | Freshwater | SAMN32031843 | NG-24277 | 5002746 | 31221 | 95.8 C:90.8%[S:87.5%,D:3.3%],F:5.0%,M:4.2% | 52452 | 92.2 C:86.3%[S:83.9%,D:2.4%],F:5.9%,M:7.8%,n:255 |
| 14 Apusomonadida | *Podomonas magna* | PodMag | JJP-2003 CCAP1901/4 | Marine | SAMN32031844 | NG-22350 | 19652204 | 93321 | 96.4 C:95.4%[S:86.5%,D:8.9%],F:1.0%,M:3.6% | 140695 | 95.3 C:92.2%[S:87.5%,D:4.7%],F:3.1%,M:4.7%,n:255 |
| 14 Apusomonadida | *Podomonas capensis* | PodCap | SPRINTER | Marine | SAMN32031845 | NG-24277 | 5028944 | 51329 | 96.4 C:94.1%[S:89.1%,D:5.0%],F:2.3%,M:3.6% | 85954 | 93.7 C:85.5%[S:83.5%,D:2.0%],F:8.2%,M:6.3%,n:255 |
| 14 Apusomonadida | *Multimonas media* | MulMed | MMROSKO2018 | Marine | SAMN32031846 | NG-24277 | 5365203 | 39146 | 93.7 C:86.8%[S:68.6%,D:18.2%],F:6.9%,M:6.3% | 97275 | 90.2 C:81.6%[S:70.6%,D:11.0%],F:8.6%,M:9.8%,n:255 |
| 14 Apusomonadida | *Apusomonas australiensis* | ApuAus | PROMEX | Freshwater | SAMN32031847 | NG-22350 | 17927843 | 45197 | 98.4 C:97.4%[S:73.3%,D:24.1%],F:1.0%,M:1.6% | 112932 | 97.2 C:95.2%[S:82.7%,D:12.5%],F:2.0%,M:2.8%,n:255 |
| 14 Apusomonadida | *Apusomonas proboscidea* | ApuPro | MPSANABRIA15 | Freshwater | SAMN32031848 | NG-24277 | 6170697 | 85522 | 97.4 C:90.1%[S:47.9%,D:42.2%],F:7.3%,M:2.6% | 119453 | 93 C:86.3%[S:47.1%,D:39.2%],F:6.7%,M:7.0%,n:255 |
| 14 Apusomonadida | *Karpovia croatica* | KarCro | CRO19P6 | Marine | SAMN32031849 | NG-24277 | 6682786 | 17839 | 95.7 C:93.1%[S:84.8%,D:8.3%],F:2.6%,M:4.3% | 66469 | 93.3 C:86.2%[S:83.1%,D:3.1%],F:7.1%,M:6.7%,n:255 |
| 14 Apusomonadida | *Singekia montserratensis* | SinMon | MONTSE19P8 | Freshwater | SAMN32031850 | NG-24277 | 5560582 | 19677 | 95.1 C:91.8%[S:87.8%,D:4.0%],F:3.3%,M:4.9% | 65350 | 93.8 C:87.1%[S:86.3%,D:0.8%],F:6.7%,M:6.2%,n:255 |
| 14 Apusomonadida | *Singekia franciliensis* | SinFra | ORSAYFEB19APU1 | Freshwater | SAMN32031851 | NG-24277 | 7738300 | 25309 | 93.4 C:88.4%[S:77.2%,D:11.2%],F:5.0%,M:6.6% | 76673 | 92.5 C:84.3%[S:80.0%,D:4.3%],F:8.2%,M:7.5%,n:255 |
| 14 Apusomonadida | *Chelonemonas dolani* | CheDol | D513M | Marine | SAMN32031852 | NG-22350 | 17648679 | 32217 | 95.7 C:93.1%[S:85.8%,D:7.3%],F:2.6%,M:4.3% | 83506 | 93.4 C:89.5%[S:87.5%,D:2.0%],F:3.9%,M:6.6%,n:255 |
| 14 Apusomonadida | *Chelonemonas geobuk* | CheGeo | KM053 | Marine | SAMN32031853 | NG-24277 | 5642089 | 18848 | 95.1 C:87.8%[S:86.5%,D:1.3%],F:7.3%,M:4.9% | 71789 | 94.1 C:85.9%[S:85.5%,D:0.4%],F:8.2%,M:5.9%,n:255 |
| 7 Ancyromonadida | *Nutomonas limna terrestris* | NutLim | ORSAYFEB19ANCY | Freshwater | SAMN32031854 | NG-24277 | 5921752 | 48297 | 80.2 C:55.1%[S:47.5%,D:7.6%],F:25.1%,M:19.8% | 70576 | 64.3 C:36.1%[S:31.0%,D:5.1%],F:28.2%,M:35.7%,n:255 |
| 7 Ancyromonadida | *Caraotamonas croatica* | CarCro | CRO19S5 | Marine | SAMN32031855 | NG-24277 | 6157385 | 30795 | 94 C:88.1%[S:83.5%,D:4.6%],F:5.9%,M:6.0% | 59629 | 87.4 C:79.2%[S:78.0%,D:1.2%],F:8.2%,M:12.6%,n:255 |
| 7 Ancyromonadida | *Ancyromonas mediterranea* | AncMed | C362 | Marine | SAMN32031856 | NG-22350 | 34510209 | 26730 | 94.1 C:87.8%[S:82.8%,D:5.0%],F:6.3%,M:5.9% | 79704 | 90.2 C:80.8%[S:78.8%,D:2.0%],F:9.4%,M:9.8%,n:255 |
| 7 Ancyromonadida | *Ancyromonas kenti* | AncKen | AKROSKO2018 | Marine | SAMN32031857 | NG-22350 | 20255134 | 116461 | 93.1 C:85.2%[S:76.9%,D:8.3%],F:7.9%,M:6.9% | 89793 | 87.1 C:75.7%[S:71.4%,D:4.3%],F:11.4%,M:12.9%,n:255 |
| 7 Ancyromonadida | *Planomonas micra* | PlaMic | PMROSKO2018 | Marine | SAMN32031858 | NG-24277 | 6485357 | 22389 | 94.6 C:84.4%[S:78.5%,D:5.9%],F:10.2%,M:5.4% | 71456 | 89.5 C:77.3%[S:75.3%,D:2.0%],F:12.2%,M:10.5%,n:255 |
| 7 Ancyromonadida | *Nyramonas silfraensis* | NyrSil | ORSLANDS2 | Freshwater | SAMN32031859 | NG-22350 | 23609535 | 28865 | 93.4 C:89.1%[S:82.2%,D:6.9%],F:4.3%,M:6.6% | 65275 | 90.3 C:83.2%[S:81.6%,D:1.6%],F:7.1%,M:9.7%,n:255 |
| 7 Ancyromonadida | *Fabomonas mesopelagica* | FabMes | A153 | Marine | SAMN32031860 | NG-25209 | 7208640 | 15724 | 98 C:94.4%[S:89.1%,D:5.3%],F:3.6%,M:2.0% | 38944 | 95.8 C:89.1%[S:87.5%,D:1.6%],F:6.7%,M:4.2%,n:255 |
| | *Meteora sporadica* | MetSpo | CRO19MET | Marine | SAMN32031861 | NG-24277 | 5936635 | 19914 | 96.4 C:94.4%[S:86.1%,D:8.3%],F:2.0%,M:3.6% | 59712 | 95.3 C:92.2%[S:90.2%,D:2.0%],F:3.1%,M:4.7%,n:255 |

| Group | Organism | CroCo: without cross-contamination – transcripts: No. transcripts | CroCo: without cross-contamination – predicted peptides + cdhit1: No. Oligopeptides | CroCo: without cross-contamination – predicted peptides + cdhit1: BUSCO |
|---|---|---|---|---|
| 14 Apusomonadida | *Mylnikovia oxoniensis* | 77325 | 149753 | 98 C:96.4%[S:32.9%,D:63.5%],F:1.6%,M:2.0%,n:255 |
| 14 Apusomonadida | *Cavaliersmithia chaoae* | 36644 | 64621 | 95.7 C:91.4%[S:85.5%,D:5.9%],F:4.3%,M:4.3%,n:255 |
| 14 Apusomonadida | *Catacumbia lutetiensis* APU2 | 38541 | 57173 | 93.7 C:84.7%[S:82.0%,D:2.7%],F:9.0%,M:6.3%,n:255 |
| 14 Apusomonadida | *Catacumbia lutetiensis* CAT | 30801 | 52271 | 92.1 C:86.2%[S:83.5%,D:2.7%],F:5.9%,M:7.9%,n:255 |
| 14 Apusomonadida | *Podomonas magna* | 89615 | 133651 | 95.3 C:92.2%[S:85.9%,D:6.3%],F:3.1%,M:4.7%,n:255 |
| 14 Apusomonadida | *Podomonas capensis* | 51242 | 85916 | 93.7 C:85.5%[S:81.2%,D:4.3%],F:8.2%,M:6.3%,n:255 |
| 14 Apusomonadida | *Multimonas media* | 38437 | 96527 | 90.2 C:81.6%[S:67.1%,D:14.5%],F:8.6%,M:9.8%,n:255 |
| 14 Apusomonadida | *Apusomonas australiensis* | 42930 | 109758 | 97.3 C:95.3%[S:80.0%,D:15.3%],F:2.0%,M:2.7%,n:255 |
| 14 Apusomonadida | *Apusomonas proboscidea* | 69257 | 100646 | 90.9 C:75.6%[S:58.0%,D:17.6%],F:15.3%,M:9.1%,n:255 |
| 14 Apusomonadida | *Karpovia croatica* | 17725 | 66418 | 93.4 C:86.3%[S:81.2%,D:5.1%],F:7.1%,M:6.6%,n:255 |
| 14 Apusomonadida | *Singekia montserratensis* | 19525 | 65227 | 93.7 C:87.0%[S:83.9%,D:3.1%],F:6.7%,M:6.3%,n:255 |
| 14 Apusomonadida | *Singekia franciliensis* | 25111 | 76529 | 92.6 C:84.4%[S:76.9%,D:7.5%],F:8.2%,M:7.4%,n:255 |
| 14 Apusomonadida | *Chelonemonas dolani* | 28911 | 78221 | 93.3 C:89.4%[S:84.7%,D:4.7%],F:3.9%,M:6.7%,n:255 |
| 14 Apusomonadida | *Chelonemonas geobuk* | 18799 | 71770 | 94.1 C:85.9%[S:85.1%,D:0.8%],F:8.2%,M:5.9%,n:255 |
| 7 Ancyromonadida | *Nutomonas limna terrestris* | 48096 | 70466 | 64.3 C:36.1%[S:29.4%,D:6.7%],F:28.2%,M:35.7%,n:255 |
| 7 Ancyromonadida | *Caraotamonas croatica* | 30189 | 59191 | 87.5 C:79.3%[S:77.3%,D:2.0%],F:8.2%,M:12.5%,n:255 |
| 7 Ancyromonadida | *Ancyromonas mediterranea* | 23582 | 74756 | 90.2 C:80.8%[S:77.3%,D:3.5%],F:9.4%,M:9.8%,n:255 |
| 7 Ancyromonadida | *Ancyromonas kenti* | 113371 | 84015 | 87.1 C:75.7%[S:68.6%,D:7.1%],F:11.4%,M:12.9%,n:255 |
| 7 Ancyromonadida | *Planomonas micra* | 21406 | 70262 | 89.3 C:76.8%[S:72.5%,D:4.3%],F:12.5%,M:10.7%,n:255 |
| 7 Ancyromonadida | *Nyramonas silfraensis* | 25546 | 59033 | 89.9 C:83.2%[S:77.3%,D:5.9%],F:6.7%,M:10.1%,n:255 |
| 7 Ancyromonadida | *Fabomonas mesopelagica* | 15703 | 38937 | 95.8 C:89.1%[S:87.5%,D:1.6%],F:6.7%,M:4.2%,n:255 |
| | *Meteora sporadica* | 19616 | 59553 | 95.2 C:92.1%[S:87.8%,D:4.3%],F:3.1%,M:4.8%,n:255 |

| Group | Organism | Diamond BAUVdb – eukaryote hits: No. oligopeptides | Diamond BAUVdb – eukaryote hits: BUSCO | Diamond BAUVdb – eukaryote hits: No. eggNOG | Diamond BAUVdb – eukaryote hits: BUSCO | Redundancy – cd-hit 90% identity: No. oligopeptides | Redundancy – cd-hit 90% identity: BUSCO |
|---|---|---|---|---|---|---|---|
| 14 Apusomonadida | *Mylnikovia oxoniensis* | 19785 | 82.4 C:81.2%[S:49.0%,D:32.2%],F:1.2%,M:17.6%,n:25[illegible] | 9651 | 75.4 C:73.4%[S:47.1%,D:26.3%],F:2.0%,M:24.6%,n:255 | 11455 | 82.4 C:81.2%[S:76.9%,D:4.3%],F:1.2%,M:17.6%,n:255 |
| 14 Apusomonadida | *Cavaliersmithia chaoae* | 10130 | 90.6 C:87.5%[S:87.1%,D:0.4%],F:3.1%,M:9.4%,n:255 | 5191 | 79.7 C:77.3%[S:76.5%,D:0.8%],F:2.4%,M:20.3%,n:255 | 9161 | 90.6 C:87.5%[S:87.5%,D:0.0%],F:3.1%,M:9.4%,n:255 |
| 14 Apusomonadida | *Catacumbia lutetiensis* APU2 | 8090 | 89.4 C:80.8%[S:79.2%,D:1.6%],F:8.6%,M:10.6%,n:255 | 4392 | 80.1 C:73.0%[S:71.0%,D:2.0%],F:7.1%,M:19.9%,n:255 | 7905 | 89.4 C:80.8%[S:80.0%,D:0.8%],F:8.6%,M:10.6%,n:255 |
| 14 Apusomonadida | *Catacumbia lutetiensis* CAT | 7923 | 88.3 C:82.4%[S:80.0%,D:2.4%],F:5.9%,M:11.7%,n:255 | 4286 | 77.3 C:72.2%[S:70.2%,D:2.0%],F:5.1%,M:22.7%,n:255 | 7713 | 88.3 C:82.4%[S:81.6%,D:0.8%],F:5.9%,M:11.7%,n:255 |
| 14 Apusomonadida | *Podomonas magna* | 12381 | 88.6 C:85.5%[S:82.0%,D:3.5%],F:3.1%,M:11.4%,n:255 | 6764 | 79.1 C:76.0%[S:72.5%,D:3.5%],F:3.1%,M:20.9%,n:255 | 11559 | 88.6 C:85.5%[S:84.7%,D:0.8%],F:3.1%,M:11.4%,n:255 |
| 14 Apusomonadida | *Podomonas capensis* | 11266 | 89.1 C:81.6%[S:80.0%,D:1.6%],F:7.5%,M:10.9%,n:255 | 6364 | 79.3 C:73.4%[S:71.8%,D:1.6%],F:5.9%,M:20.7%,n:255 | 10692 | 89.1 C:81.6%[S:80.4%,D:1.2%],F:7.5%,M:10.9%,n:255 |
| 14 Apusomonadida | *Multimonas media* | 11967 | 84.6 C:76.8%[S:68.2%,D:8.6%],F:7.8%,M:15.4%,n:255 | 6040 | 74.2 C:66.7%[S:59.6%,D:7.1%],F:7.5%,M:25.8%,n:255 | 9513 | 84.7 C:76.9%[S:75.7%,D:1.2%],F:7.8%,M:15.3%,n:255 |
| 14 Apusomonadida | *Apusomonas australiensis* | 14430 | 90.6 C:89.0%[S:77.6%,D:11.4%],F:1.6%,M:9.4%,n:255 | 8248 | 81.6 C:80.4%[S:70.2%,D:10.2%],F:1.2%,M:18.4%,n:255 | 12194 | 90.6 C:89.0%[S:83.9%,D:5.1%],F:1.6%,M:9.4%,n:255 |
| 14 Apusomonadida | *Apusomonas proboscidea* | 9192 | 63.9 C:49.0%[S:43.5%,D:5.5%],F:14.9%,M:36.1%,n:25[illegible] | 5367 | 59.2 C:46.3%[S:40.8%,D:5.5%],F:12.9%,M:40.8%,n:255 | 9076 | 63.9 C:49.0%[S:43.5%,D:5.5%],F:14.9%,M:36.1%,n:255 |
| 14 Apusomonadida | *Karpovia croatica* | 9291 | 88.2 C:81.5%[S:78.8%,D:2.7%],F:6.7%,M:11.8%,n:255 | 5083 | 77.6 C:71.7%[S:69.0%,D:2.7%],F:5.9%,M:22.4%,n:255 | 9096 | 88.2 C:81.5%[S:78.8%,D:2.7%],F:6.7%,M:11.8%,n:255 |
| 14 Apusomonadida | *Singekia montserratensis* | 9075 | 89.4 C:83.9%[S:83.1%,D:0.8%],F:5.5%,M:10.6%,n:255 | 4733 | 74.6 C:72.2%[S:71.4%,D:0.8%],F:2.4%,M:25.4%,n:255 | 8943 | 89.4 C:83.9%[S:83.1%,D:0.8%],F:5.5%,M:10.6%,n:255 |
| 14 Apusomonadida | *Singekia franciliensis* | 9101 | 89 C:80.4%[S:78.0%,D:2.4%],F:8.6%,M:11.0%,n:255 | 4986 | 74.6 C:67.1%[S:64.7%,D:2.4%],F:7.5%,M:25.4%,n:255 | 8685 | 89 C:80.4%[S:78.8%,D:1.6%],F:8.6%,M:11.0%,n:255 |
| 14 Apusomonadida | *Chelonemonas dolani* | 11352 | 92.9 C:89.0%[S:88.6%,D:0.4%],F:3.9%,M:7.1%,n:255 | 7046 | 87.8 C:84.3%[S:84.3%,D:0.0%],F:3.5%,M:12.2%,n:255 | 11128 | 92.9 C:89.0%[S:89.0%,D:0.0%],F:3.9%,M:7.1%,n:255 |
| 14 Apusomonadida | *Chelonemonas geobuk* | 12141 | 92.9 C:84.7%[S:84.3%,D:0.4%],F:8.2%,M:7.1%,n:255 | 7181 | 89 C:81.2%[S:81.2%,D:0.0%],F:7.8%,M:11.0%,n:255 | 11963 | 92.9 C:84.7%[S:84.3%,D:0.4%],F:8.2%,M:7.1%,n:255 |
| 7 Ancyromonadida | *Nutomonas limna terrestris* | 5514 | 55.7 C:30.2%[S:27.1%,D:3.1%],F:25.5%,M:44.3%,n:25[illegible] | 3294 | 49.1 C:26.7%[S:24.3%,D:2.4%],F:22.4%,M:50.9%,n:255 | 5322 | 55.7 C:30.2%[S:28.2%,D:2.0%],F:25.5%,M:44.3%,n:255 |
| 7 Ancyromonadida | *Caraotamonas croatica* | 5552 | 80.4 C:72.6%[S:71.8%,D:0.8%],F:7.8%,M:19.6%,n:255 | 2938 | 65.1 C:59.2%[S:58.0%,D:1.2%],F:5.9%,M:34.9%,n:255 | 5491 | 80.4 C:72.6%[S:71.8%,D:0.8%],F:7.8%,M:19.6%,n:255 |
| 7 Ancyromonadida | *Ancyromonas mediterranea* | 5807 | 80.9 C:73.4%[S:71.8%,D:1.6%],F:7.5%,M:19.1%,n:255 | 3156 | 65.5 C:59.6%[S:57.6%,D:2.0%],F:5.9%,M:34.5%,n:255 | 5569 | 80.9 C:73.4%[S:72.2%,D:1.2%],F:7.5%,M:19.1%,n:255 |
| 7 Ancyromonadida | *Ancyromonas kenti* | 5583 | 78.1 C:68.7%[S:66.3%,D:2.4%],F:9.4%,M:21.9%,n:255 | 3029 | 64 C:56.9%[S:54.5%,D:2.4%],F:7.1%,M:36.0%,n:255 | 5308 | 77.6 C:68.6%[S:68.2%,D:0.4%],F:9.0%,M:22.4%,n:255 |
| 7 Ancyromonadida | *Planomonas micra* | 6065 | 82 C:72.2%[S:71.4%,D:0.8%],F:9.8%,M:18.0%,n:255 | 3609 | 71.7 C:63.9%[S:63.1%,D:0.8%],F:7.8%,M:28.3%,n:255 | 5953 | 82 C:72.2%[S:71.8%,D:0.4%],F:9.8%,M:18.0%,n:255 |
| 7 Ancyromonadida | *Nyramonas silfraensis* | 5764 | 81.2 C:76.5%[S:75.3%,D:1.2%],F:4.7%,M:18.8%,n:255 | 3213 | 68.6 C:65.5%[S:64.3%,D:1.2%],F:3.1%,M:31.4%,n:255 | 5577 | 81.2 C:76.5%[S:75.3%,D:1.2%],F:4.7%,M:18.8%,n:255 |
| 7 Ancyromonadida | *Fabomonas mesopelagica* | 6662 | 89.1 C:82.8%[S:81.6%,D:1.2%],F:6.3%,M:10.9%,n:255 | 3885 | 77.7 C:73.0%[S:71.4%,D:1.6%],F:4.7%,M:22.3%,n:255 | 6519 | 89.1 C:82.8%[S:82.0%,D:0.8%],F:6.3%,M:10.9%,n:255 |
| | *Meteora sporadica* | 5197 | 87.5 C:84.4%[S:82.4%,D:2.0%],F:3.1%,M:12.5%,n:255 | 3156 | 74.6 C:73.4%[S:71.8%,D:1.6%],F:1.2%,M:25.4%,n:255 | 4964 | 87.4 C:84.3%[S:83.5%,D:0.8%],F:3.1%,M:12.6%,n:255 |

**Table S3. Cross-contamination summary for each RNA sequencing batch.** Data were parsed from CroCo network's LINKS_diagrammer output files. Taxa in bold are used in this study (the rest correspond to sequences from other

| batch NG-22350 | **AKROSKO2018** | **C362** | **ORSLAND19S2** | CCMP17 | CCMP18 | CCMP26 | E3stram | Guifr | NY0222 | NY0223 | NY0224 | **D513M** | **PROMEX** | **Pmagna** | **Toxo** | Total |
|---|---|---|---|---|---|---|---|---|---|---|---|---|---|---|---|---|
| **AKROSKO2018** | 0 | 26 | 30 | 94 | 28 | 50 | 68 | 93 | 29 | 41 | 18 | 58 | 44 | 57 | 21 | 657 |
| **C362** | 171 | 0 | 42 | 79 | 44 | 47 | 55 | 87 | 38 | 38 | 25 | 270 | 41 | 59 | 31 | 1027 |
| **ORSLAND19S2** | 105 | 26 | 0 | 112 | 37 | 58 | 76 | 115 | 37 | 32 | 28 | 85 | 51 | 56 | 44 | 862 |
| CCMP17 | 36 | 2 | 11 | 0 | 117 | 20 | 18 | 57 | 8 | 14 | 2 | 26 | 6 | 26 | 10 | 353 |
| CCMP18 | 45 | 11 | 25 | 70 | 0 | 33 | 36 | 77 | 27 | 40 | 9 | 41 | 34 | 43 | 18 | 509 |
| CCMP26 | 22 | 7 | 22 | 109 | 103 | 0 | 22 | 48 | 18 | 15 | 5 | 42 | 21 | 24 | 9 | 467 |
| E3stram | 83 | 22 | 28 | 77 | 32 | 48 | 0 | 83 | 27 | 45 | 20 | 62 | 25 | 60 | 16 | 628 |
| Guifr | 39 | 18 | 37 | 102 | 115 | 50 | 46 | 0 | 33 | 21 | 13 | 53 | 31 | 44 | 16 | 618 |
| NY0222 | 29 | 15 | 24 | 40 | 17 | 26 | 30 | 35 | 0 | 27 | 7 | 38 | 22 | 20 | 14 | 344 |
| NY0223 | 38 | 17 | 19 | 106 | 63 | 39 | 20 | 66 | 17 | 0 | 16 | 33 | 24 | 29 | 18 | 505 |
| NY0224 | 9 | 5 | 5 | 16 | 5 | 4 | 9 | 9 | 3 | 3 | 0 | 13 | 6 | 12 | 1 | 100 |
| **D513M** | 47 | 671 | 33 | 62 | 31 | 35 | 45 | 78 | 32 | 30 | 15 | 0 | 36 | 48 | 22 | 1185 |
| **PROMEX** | 50 | 26 | 35 | 67 | 32 | 41 | 30 | 64 | 25 | 37 | 21 | 68 | 0 | 32 | 36 | 564 |
| **Pmagna** | 98 | 21 | 52 | 136 | 40 | 57 | 56 | 135 | 27 | 53 | 24 | 89 | 47 | 0 | 31 | 866 |
| **Toxo** | 46 | 13 | 36 | 53 | 36 | 25 | 25 | 66 | 13 | 22 | 11 | 44 | 33 | 23 | 0 | 446 |

| batch NG-24277 | **CRO19S5** | **ORSAYFEB19ANCY** | **PMROSKO2018** | CRO19MAN | **CRO19MET** | **CRO19P6** | **CAT** | **FABANU** | **KM053** | **MMROSKO2018** | **MONTSE19P8** | **MPSANABRIA15** | **MPcont** | **ORSAYFEB19APU1** | **ORSAYFEB19APU2** | **SPRINTER** | Total |
|---|---|---|---|---|---|---|---|---|---|---|---|---|---|---|---|---|---|
| **CRO19S5** | 0 | 4 | 18 | 8 | 35 | 3 | 0 | 23 | 4 | 8 | 0 | 0 | 1 | 0 | 0 | 2 | 106 |
| **ORSAYFEB19ANCY** | 2 | 0 | 2 | 12 | 2 | 0 | 1 | 0 | 0 | 0 | 1 | 0 | 1 | 13 | 1 | 1 | 36 |
| **PMROSKO2018** | 108 | 5 | 0 | 2 | 0 | 0 | 0 | 45 | 6 | 18 | 0 | 0 | 0 | 0 | 1 | 2 | 187 |
| CRO19MAN | 7 | 6 | 3 | 0 | 3 | 18 | 0 | 3 | 3 | 1 | 2 | 0 | 0 | 1 | 2 | 0 | 49 |
| **CRO19MET** | 15 | 7 | 0 | 113 | 0 | 2 | 0 | 0 | 5 | 1 | 1 | 0 | 0 | 0 | 1 | 0 | 145 |
| **CRO19P6** | 2 | 2 | 1 | 9 | 3 | 0 | 0 | 1 | 2 | 0 | 0 | 0 | 0 | 0 | 0 | 2 | 22 |
| **CAT** | 1 | 30 | 1 | 2 | 1 | 1 | 0 | 1 | 0 | 0 | 2 | 0 | 1 | 7 | 11 | 9 | 67 |
| **FABANU** | 28 | 4 | 6 | 2 | 43 | 1 | 2 | 0 | 2 | 12 | 0 | 1 | 2 | 0 | 0 | 2 | 105 |
| **KM053** | 0 | 2 | 6 | 1 | 0 | 0 | 2 | 2 | 0 | 2 | 0 | 0 | 1 | 0 | 1 | 1 | 18 |
| **MMROSKO2018** | 144 | 2 | 26 | 4 | 0 | 2 | 0 | 18 | 1 | 0 | 0 | 0 | 2 | 0 | 0 | 0 | 199 |
| **MONTSE19P8** | 1 | 13 | 0 | 0 | 0 | 0 | 1 | 0 | 0 | 0 | 0 | 0 | 2 | 36 | 1 | 0 | 54 |
| **MPSANABRIA15** | 3 | 16 | 0 | 6 | 1 | 1 | 3 | 3 | 0 | 1 | 7 | 0 | 2398 | 2 | 1 | 4 | 2446 |
| **MPcontaminant** | 0 | 18 | 0 | 9 | 0 | 0 | 8 | 2 | 1 | 0 | 10 | 342 | 0 | 3 | 3 | 3 | 399 |
| **ORSAYFEB19APU1** | 1 | 30 | 0 | 5 | 1 | 0 | 1 | 1 | 1 | 1 | 3 | 0 | 2 | 0 | 0 | 0 | 46 |
| **ORSAYFEB19APU2** | 0 | 56 | 1 | 8 | 3 | 0 | 40 | 4 | 0 | 1 | 2 | 0 | 2 | 11 | 0 | 1 | 129 |
| **SPRINTER** | 4 | 3 | 2 | 3 | 0 | 0 | 1 | 1 | 3 | 0 | 2 | 0 | 0 | 0 | 2 | 0 | 21 |

| batch NG-25209 | **A153** | aphO14 | aphP2 | CRO20ST1 | ORSAY19EX4 | Total |
|---|---|---|---|---|---|---|
| **A153** | 0 | 0 | 1 | 8 | 2 | 11 |
| aphO14 | 1 | 0 | 3776 | 23 | 26 | 3826 |
| aphP2 | 3 | 4193 | 0 | 26 | 29 | 4251 |
| CRO20ST1 | 3 | 3 | 1 | 0 | 29 | 36 |
| ORSAY19EX4 | 2 | 6 | 4 | 26 | 0 | 38 |

Table S4. List of 32 eukaryotes with reference genomes used to extract eukaryotic peptides from the metatranscriptomes.

| Species name | Version |
|---|---|
| *Homo sapiens* | Ensembl 102 |
| *Aplysia californica* | NCBI GCF_000002075.1_AplCal3.0 |
| *Salpingoeca rosetta* | Ensembl Protists 49 |
| *Capsaspora owczarzaki* | Ensembl Protists 49 |
| *Sphaeroforma arctica* | Dudin et al. 2019 (Sarc4), Figshare: https://figshare.com/articles/Sphaeroforma_arctica_transcriptome/8299529 |
| *Corallochytrium limacisporum* | Grau-Bové et al. 2017, Figshare: https://figshare.com/articles/Genome_-_Corallochytrium_limacisporum/5426470 |
| *Neurospora crassa* | Ensembl Fungi 49, NC12 |
| *Cryptococcus neoformans* | Ensembl Fungi 49, ASM9104v1 (chromosomal assembly) |
| *Spizellomyces punctatus* | Ensembl Fungi 49, V1, DAOM BR117 |
| *Rozella allomycis* | Ensembl Fungi 49, Rozella_k41_t100, CSF55 |
| *Fonticula alba* | NCBI GCF_000388065.1; assembly Font_alba_ATCC_38817_V2 |
| *Thecamonas trahens* | Ensembl Protist 39, v1 2010, ATCC 50062 |
| *Dictyostelium discoideum* | Ensembl Protist 49, dicty_2.7 |
| *Acanthamoeba castellanii* | Ensembl Protist 49, strain Neff gca_000313135 (v1, better than so-called v2 GCA_000193105.1 in NCBI) |
| *Arabidopsis thaliana* | Ensembl Plants 49, TAIR10 |
| *Volvox carteri* | NCBI GCF_000143455.1_v1.0 |
| *Porphyridium purpureum* | NCBI GCA_008690995.1_P_purpureum_CCMP1328_Hybrid_assembly |
| *Guillardia theta* | Ensembl Protists 49, Guith1, CCMP2712 |
| *Emiliania huxleyi* | Ensembl Protists 49, CCMP1516 |
| *Ectocarpus siliculosus* | Ensembl Protists 49, gca_000310025.ASM31002v1, chromosomal assembly |
| *Phaeodactylum tricornutum* | Ensembl Protists 49, ASM15095v2 |
| *Phytophthora infestans* | Ensembl Protists 49, ASM14294v1 |
| *Hondaea fermentalgiana* | Ensembl Protists 49, gca_002897355.Aurantiochytrium_FCC1311_v1 |
| *Plasmodium falciparum* | Ensembl Protists 49 ASM276v2 3D7 |
| *Chromera velia* | CryptoDB CCMP2878 https://cryptodb.org/common/downloads/Current_Release/CveliaCCMP2878 |
| *Symbiodinium microadriaticum* | Ensembl Protists 49, gca_001939145.ASM193914v1 |
| *Paramecium tetraurelia* | Ensembl Protists 49, ASM16542v1; need to manually fix transcripts, ciliate code! |
| *Reticulomyxa filosa* | Ensembl Protists 49 gca_000512085 |
| *Bigelowiella natans* | Ensembl Protists 49, Bigna1 |
| *Naegleria gruberi* | Ensembl Protists 49, gca_000004985.V1.0 |
| *Trichomonas vaginalis* | NCBI GCF_000002825.2_ASM282v1, G3 strain |
| *Kipferlia bialata* | Ensembl Protists 49, gca_003568945 |

Table S5. Species and dataset used for the phylogenomic analysis of eukaryotes. In total, 101 species and 303 protein markers were used. Species for which the transcriptome was generated in this study are highlighted in bold.

| Species name | Name in dataset | Supergroup | No. of markers | No. missing markers | % missing markers | % Gaps | Source |
|---|---|---|---|---|---|---|---|
| *Digyalum oweni* WS1+WS2 | ALV_Dig_owe | Alveolata | 214 | 89 | 29.37 | 29.13 | Janouškovec et al. 2019 |
| *Oxytricha trifallax* | ALV_Oxy_tri | Alveolata | 280 | 23 | 7.59 | 11.62 | Swart et al. 2013 |
| *Colponema vietnamica* Colp-7a | COL_Col_vie | Alveolata | 230 | 73 | 24.09 | 27.34 | Tikhonenkov et al. 2020 |
| Colponemid sp. Colp-10 | COL_Colp10 | Alveolata | 275 | 28 | 9.24 | 7.17 | Tikhonenkov et al. 2020 |
| *Vitrella brassicaformis* CCMP3155 | VitrBras | Alveolata | 237 | 66 | 21.78 | 23.30 | Lax et al. 2018 |
| *Acanthamoeba castellanii* Neff | AcanGEN | Amoebozoa | 242 | 61 | 20.13 | 23.98 | amoebadb.org |
| *Mastigamoeba balamuthi* | AMO_Mas_bal | Amoebozoa | 236 | 67 | 22.11 | 40.64 | Nývltová et al. 2013 |
| *Phalansterium solitarum* TexasCypress | AMO_Pha_sol | Amoebozoa | 290 | 13 | 4.29 | 11.01 | Kang et al. 2017 |
| *Stenamoeba stenopodia* CCAP 1565/3 | AMO_Ste_ste | Amoebozoa | 281 | 22 | 7.26 | 11.98 | Kang et al. 2017 |
| *Vermamoeba vermiformis* ATCC 50236 | AMO_ver_ver | Amoebozoa | 176 | 127 | 41.91 | 70.10 | Bullerwell et al. 2010 |
| *Physarum polycephalum* LU352 | PhysTSA | Amoebozoa | 283 | 20 | 6.60 | 11.01 | Lax et al. 2018 |
| ***Ancyromonas mediterranea* C362** | ANC_Anc_C362_brut | Ancyromonadida | 252 | 51 | 16.83 | 14.93 | This study |
| ***Caraotamonas croatica* CRO19S5** | ANC_Anc_CRO19S5_brut | Ancyromonadida | 288 | 15 | 4.95 | 8.56 | This study |
| ***Ancyromonas kenti* AKROSKO2018** | ANC_Anc_ken_brut | Ancyromonadida | 283 | 20 | 6.60 | 9.50 | This study |
| ***Fabomonas mesopelagica* A153** | ANC_Fab_tro_A153_bru | Ancyromonadida | 291 | 12 | 3.96 | 6.73 | This study |
| ***Nutomonas limna terrestris* ORSAYFEB19ANCY** | ANC_Nuthow_brut | Ancyromonadida | 260 | 43 | 14.19 | 24.37 | This study |
| ***Nyramonas silfraensis* ORSLANDS2** | ANC_ORSLAND19S2_brut | Ancyromonadida | 291 | 12 | 3.96 | 7.97 | This study |
| ***Planomonas micra* PMROSKO2018** | ANC_Pla_mic_brut | Ancyromonadida | 286 | 17 | 5.61 | 9.88 | This study |
| *Ancyromonas sigmoides* CCAP 1958/3 (= B-70) | AncB70 | Ancyromonadida | 222 | 81 | 26.73 | 42.83 | Brown et al. 2018 |
| *Striomonas longa* CCAP 1958/5 (formerly *Nutomonas* ) | Nutolong | Ancyromonadida | 258 | 45 | 14.85 | 15.98 | Torruella et al. 2015 (uncontaminated in figshare) |
| *Fabomonas tropica* NYK3C | NYK3C | Ancyromonadida | 265 | 38 | 12.54 | 24.17 | Brown et al. 2018 |
| *Thecamonas trahens* ATCC 50062 | Thectrah | Apusomonadida | 271 | 32 | 10.56 | 13.10 | Torruella et al. 2012 |
| *Manchomonas bermudensis* ATCC 50234 (formerly *Amastigomonas* ) | APU_Man_ber | Apusomonadida | 62 | 241 | 79.54 | 90.49 | Brown et al. 2013 |
| *Singekia* sp. FB-2015 | APU_Ama_FB15 | Apusomonadida | 261 | 42 | 13.86 | 25.98 | Burki et al. 2016 |
| ***Singekia franciliensis* ORSAYFEB19APU1** | APU_APU1_brut | Apusomonadida | 290 | 13 | 4.29 | 6.64 | This study |
| ***Catacumbia lutetiensis* ORSAYFEB19APU2** | APU_APU2_brut | Apusomonadida | 297 | 6 | 1.98 | 4.80 | This study |
| ***Apusomonas australiensis* PROMEX** | APU_Apu_mex_brut | Apusomonadida | 295 | 8 | 2.64 | 5.28 | This study |
| ***Apusomonas proboscidea* MPSANABRIA15** | APU_Apu_MP15_brut | Apusomonadida | 222 | 81 | 26.73 | 29.24 | This study |
| ***Catacumbia lutetiensis* CAT** | APU_CAT_brut | Apusomonadida | 297 | 6 | 1.98 | 4.50 | This study |
| ***Chelonemonas dolani* D513M** | APU_Che_D513_brut | Apusomonadida | 253 | 50 | 16.50 | 14.52 | This study |
| ***Chelonemonas geobuk* KM053** | APU_Che_geo_brut | Apusomonadida | 291 | 12 | 3.96 | 6.39 | This study |
| ***Karpovia croatica* CRO19P6** | APU_CRO19P6_brut | Apusomonadida | 291 | 12 | 3.96 | 5.77 | This study |
| ***Cavaliersmithia chaoae* FABANU** | APU_FABANU_brut | Apusomonadida | 299 | 4 | 1.32 | 3.94 | This study |
| ***Singekia montserratensis* MONTSE19P8** | APU_MONTSE19P8_brut | Apusomonadida | 294 | 9 | 2.97 | 4.71 | This study |
| ***Multimonas media* MMROSKO2018** | APU_Mul_med_brut | Apusomonadida | 288 | 15 | 4.95 | 8.21 | This study |
| ***Podomonas capensis* SPRINTER** | APU_Pod_cap_brut | Apusomonadida | 292 | 11 | 3.63 | 5.12 | This study |
| ***Podomonas magna* CCAP 1901/4** | APU_Pod_mag_brut | Apusomonadida | 289 | 14 | 4.62 | 6.07 | This study |
| ***Mylnikovia oxoniensis* CCAP 1901/2** (formerly *Thecamonas* ) | APU_The_oxo_brut | Apusomonadida | 290 | 13 | 4.29 | 6.27 | This study |
| *Galdieria sulphuraria* | ARC_Gal_sul | Archaeplastida | 253 | 50 | 16.50 | 17.96 | Lax et al. 2018 |
| *Gloeochaete witrockiana* | ARC_Glo_wit | Archaeplastida | 270 | 33 | 10.89 | 14.22 | Lax et al. 2018 |
| *Mesostigma viride* | ARC_Mes_vir | Archaeplastida | 250 | 53 | 17.49 | 32.00 | Lax et al. 2018 |
| *Rhodelphis limneticus* | ARC_Rho_lim | Archaeplastida | 279 | 24 | 7.92 | 10.64 | Gawryluk et al. 2019 |
| *Rhodelphis marinus* | ARC_Rho_mar | Archaeplastida | 288 | 15 | 4.95 | 5.86 | Gawryluk et al. 2019 |
| *Cyanoptyche gloeocystis* SAG4.97 | CyanGloe | Archaeplastida | 215 | 88 | 29.04 | 38.81 | Lax et al. 2018 |
| *Picozoa* sp. PB584-11 | PB58411a | Archaeplastida | 93 | 210 | 69.31 | 81.82 | Lax et al. 2018 |
| *Rhodosorus marinus* | RhodMari | Archaeplastida | 226 | 77 | 25.41 | 23.56 | Lax et al. 2018 |
| *Volvox carteri* | Volvcart | Archaeplastida | 269 | 34 | 11.22 | 13.74 | Lax et al. 2018 |
| *Breviata anathema* ATCC 50338 (= *Mastigamoeba invertens* ) | BRE_Bre_ans | Breviatea | 138 | 165 | 54.46 | 72.69 | Stairs et al. 2014 |
| *Lenisia limosa* EH-2015 | BRE_Len_lim | Breviatea | 70 | 233 | 76.90 | 93.73 | Hamann et al. 2016 |
| *Subulatomonas tetraspora* ATCC 50623 | BRE_Sub_tet | Breviatea | 90 | 213 | 70.30 | 85.71 | Grant et al. 2012 |
| *Pygsuia biforma* | PCBtrin | Breviatea | 262 | 41 | 13.53 | 22.23 | Brown et al. 2013 |
| *Mantamonas plastica* CCAP 1946/1 Bass1 | Bass1 | CRuMs | 277 | 26 | 8.58 | 19.97 | Brown et al. 2018 |
| *Collodictyon triciliatum* | Colltric | CRuMs | 75 | 228 | 75.25 | 90.07 | Zhao et al. 2012 |
| *Mantamonas vickermanii* | CRO19MAN_longest_orf | CRuMs | 296 | 7 | 2.31 | 6.46 | Blaz et al. 2023 |
| *Diphylleia rotans* NIES-3764 | DiphSRT | CRuMs | 289 | 14 | 4.62 | 7.34 | Brown et al. 2018 |
| *Rigifila ramosa* CCAP 1967/1 | Rigiramo | CRuMs | 277 | 26 | 8.58 | 12.56 | Brown et al. 2018 |
| *Goniomonas pacifica* CCMP1869 | CRY_Gon_pac | Cryptista | 286 | 17 | 5.61 | 13.42 | Lax et al. 2018 |
| *Guillardia theta* CCMP2712 | CRY_Gui_the | Cryptista | 284 | 19 | 6.27 | 11.05 | Lax et al. 2018 |
| *Palpitomonas bilix* MMETSP0780 | CRY_Pal_bil | Cryptista | 250 | 53 | 17.49 | 27.07 | Lax et al. 2018 |
| *Roombia truncata* | CRY_Roo_tru | Cryptista | 215 | 88 | 29.04 | 52.76 | Lax et al. 2018 |
| Cryptophyceae sp. | Cryp2293 | Cryptista | 238 | 65 | 21.45 | 24.88 | Lax et al. 2018 |
| *Cryptomonas paramecium* | Cryppara | Cryptista | 259 | 44 | 14.52 | 21.59 | Lax et al. 2018 |
| *Andalucia godoyi* And28 | DIS_And_god | Discoba | 270 | 33 | 10.89 | 14.98 | Gray et al., 2020 |
| *Euglena gracilis* Z1 | DIS_Eug_gra | Discoba | 179 | 124 | 40.92 | 65.16 | Ebenezer et al. 2019 |
| *Pharyngomonas kirbyi* | DIS_Pha_kir | Discoba | 288 | 15 | 4.95 | 7.86 | Lax et al. 2018 |
| *Stygella incarcerata* MB1 | DIS_Sty_inc | Discoba | 118 | 185 | 61.06 | 77.70 | Leger et al. 2016 |
| *Tsukubamonas globosa* | Tsukglob | Discoba | 208 | 95 | 31.35 | 66.00 | Lax et al. 2018 |
| *Acanthocystis* sp. | HAP_Aca_sp | Haptista | 239 | 64 | 21.12 | 23.29 | Lax et al. 2018 |
| *Emiliania huxleyi* CCMP1516 | HAP_Emi_hux | Haptista | 242 | 61 | 20.13 | 28.06 | Lax et al. 2018 |
| Pavlovales sp. CCMP2436 | HAP_Pav_sp | Haptista | 241 | 62 | 20.46 | 27.12 | Lax et al. 2018 |
| *Pavlova lutheri* | Pavlova | Haptista | 129 | 174 | 57.43 | 79.02 | Lax et al. 2018 |
| *Prymnesium parvum* | Prymparv | Haptista | 255 | 48 | 15.84 | 22.13 | Lax et al. 2018 |
| *Hemimastix kukwesjijk* | Hemimast | Hemimastigophora | 240 | 63 | 20.79 | 30.87 | Lax et al. 2018 |
| *Spironema multiciliatum* | Spironem | Hemimastigophora | 249 | 54 | 17.82 | 25.19 | Lax et al. 2018 |
| *Gefionella okellyi* Mal249 | Mal249 | Malawimonadida | 260 | 43 | 14.19 | 18.66 | Heiss et al. 2018 |
| *Malawimonas jakobiformis* | MalaJako | Malawimonadida | 167 | 136 | 44.88 | 69.48 | Lax et al. 2018 |
| *Carpediemonas membranifera* | MET_Car_mem | Metamonada | 174 | 129 | 42.57 | 44.42 | Leger et al. 2017 |
| *Dysnectes brevis* | MET_Dys_bre | Metamonada | 236 | 67 | 22.11 | 30.23 | Leger et al. 2017 |
| *Trimastix marina* PCT | MET_Tri_mar | Metamonada | 225 | 78 | 25.74 | 33.39 | Leger et al. 2017 |
| *Trichomonas vaginalis* | MET_Tri_vag | Metamonada | 215 | 88 | 29.04 | 31.98 | https://trichdb.org/common/downloads/release-47/TvaginalisG3/fasta/data/ |
| *Paratrimastix pyriformis* RCP-MX ATCC 50935 | Trimpyri | Metamonada | 141 | 162 | 53.47 | 72.00 | Lax et al. 2018 |
| ***Meteora sporadica* CRO19MET** | CRO19MET_longest_orf | Meteora | 277 | 26 | 8.58 | 11.48 | This study |
| *Parvularia atlantis* ATCC 50694 | OPI_Par_atl | Opisthokonta | 278 | 25 | 8.25 | 13.21 | Torruella et al. 2015 (uncontaminated in figshare) |
| *Paraphelidium tribonematis* x-108 (= *P. tribonemae* ) | OPI_Par_tri | Opisthokonta | 279 | 24 | 7.92 | 9.88 | Torruella et al. 2018 (uncontaminated in figshare) |
| *Pigoraptor chileana* Opistho-2 | OPI_Pig_chi | Opisthokonta | 271 | 32 | 10.56 | 17.53 | Hehenberger et al. 2017 |
| *Salpingoeca rosetta* ATCC 50818 | OPI_Sal_ros | Opisthokonta | 237 | 66 | 21.78 | 26.47 | Torruella et al. 2012 |
| *Syssomonas multiformis* Colp-12 (= Opisthokonta sp. DVT-2017c) | OPI_Sys_mul | Opisthokonta | 268 | 35 | 11.55 | 22.17 | Hehenberger et al. 2017 |
| *Spizellomyces punctatus* DAOM BR117 | Spizpunc | Opisthokonta | 285 | 18 | 5.94 | 7.87 | Lax et al. 2018 |
| *Ancoracysta twista* TD-1 | HAP_Anc_twi | Provora | 284 | 19 | 6.27 | 16.28 | Janouškovec et al. 2017 |
| *Paulinella chromatophora* CCAC0185 | Paulchro | Rhizaria | 216 | 87 | 28.71 | 61.56 | Lax et al. 2018 |
| *Bigelowiella natans* CCMP2755 | RHI_Big_nat | Rhizaria | 277 | 26 | 8.58 | 14.89 | Lax et al. 2018 |
| *Lapot gusevi* Lap-1 | RHI_Lap_gus | Rhizaria | 265 | 38 | 12.54 | 15.75 | Irwin et al. 2019 |
| *Reticulomyxa filosa* | RHI_Ret_fil | Rhizaria | 251 | 52 | 17.16 | 31.03 | Glöckner et al. 2013 |
| *Spongosphaera subterranea* | RHI_Spo_sub | Rhizaria | 193 | 110 | 36.30 | 42.46 | Schwelm et al. 2015 |
| *Aplanochytrium kerguelense* | Aplakerg | Stramenopila | 274 | 29 | 9.57 | 12.07 | Lax et al. 2018 |
| *Ectocarpus siliculosus* Ec 32 CCAP 1310/04 | Ectosili | Stramenopila | 287 | 16 | 5.28 | 12.72 | Lax et al. 2018 |
| *Phytophthora infestans* | Phytinfe | Stramenopila | 277 | 26 | 8.58 | 11.04 | Lax et al. 2018 |
| *Halocafeteria seosinensis* | STR_Hal_seo | Stramenopila | 292 | 11 | 3.63 | 5.79 | Harding et al. 2016 |
| *Platysulcus tardus* | STR_Pla_tar | Stramenopila | 271 | 32 | 10.56 | 10.79 | Shiratori et al. 2015 |
| Telonemid P2 | TEL_Tel_p2 | Telonemia | 274 | 29 | 9.57 | 12.66 | Strassert et al. 2019 |
| *Telonema subtile* | TEL_Tel_sub | Telonemia | 278 | 25 | 8.25 | 13.15 | Strassert et al. 2019 |

Table S6. Alternative topology tests of phylogenomic analyses. IQ-tree output of testing constrained trees. The KH, SH and AU tests return p-values (< 0.05 for rejection).

| Constrained monophyly | Tree | logL | deltaL | bp-RELL | p-KH | p-SH | c-ELW | p-AU |
|---|---|---|---|---|---|---|---|---|
| malawimonads + ancyromonads (unconstrained) | 1 | -7293812.346 | 0 | 0.403 | 0.601 | 1 | 0.403 | 0.624 |
| malawimonads + discobans | 2 | -7293916.904 | 104.56 | 0.0036 | 0.124 | 0.0546 | 0.00366 | 0.0257 |
| malawimonads + metamonads | 3 | -7293840.378 | 28.031 | 0.1 | 0.253 | 0.53 | 0.1 | 0.237 |
| malawimonads + podiates | 4 | -7293818.61 | 6.2633 | 0.186 | 0.399 | 0.81 | 0.187 | 0.471 |
| metamonads + ancyromonads | 5 | -7293819.388 | 7.0421 | 0.307 | 0.437 | 0.714 | 0.307 | 0.501 |
